# Species-specific LUBAC-mediated M1 ubiquitination regulates necroptosis by segregating the cellular distribution and fate of activated MLKL

**DOI:** 10.1101/2022.12.08.519265

**Authors:** Nadine Weinelt, Kaja Nicole Wächtershäuser, Sonja Smith, Geoffroy Andrieux, Tonmoy Das, Birte Jeiler, Jens Roedig, Leonard Feist, Björn Rotter, Melanie Boerries, Francesco Pampaloni, Sjoerd J. L. van Wijk

## Abstract

Plasma membrane accumulation of phosphorylated mixed lineage kinase domain-like (MLKL) is a hallmark of necroptosis, leading to membrane rupture and inflammatory cell death. Pro-death functions of MLKL are tightly controlled by several checkpoints, including phosphorylation. Endocytosis and exocytosis limit MLKL membrane accumulation and counteract necroptosis, but the exact mechanisms remain poorly understood. Here, we identify linear ubiquitin chain assembly complex (LUBAC)-mediated M1 poly-ubiquitination (poly-Ub) as novel checkpoint for necroptosis regulation downstream of activated MLKL in human cells. Loss of LUBAC activity inhibits necroptosis, without affecting necroptotic signaling, but by preventing membrane accumulation of activated MLKL. Flotillin-1/2 act as putative necroptotic M1 poly-Ub targets that inhibit necroptosis suppression induced by LUBAC inhibition. Finally, we confirm LUBAC-dependent activation of necroptosis in primary human pancreatic organoids. Our findings identify LUBAC as species-specific regulator of necroptosis which promotes MLKL membrane accumulation and pioneer primary human organoids to model necroptosis in near-physiological settings.

## Introduction

Caspase-independent necroptosis, characterized by plasma membrane rupture, release of cytokines, chemokines and damage-associated molecular patterns (DAMPs), elicit potent immunogenic responses ^1, 2^. Accordingly, necroptotic cell death underlies a wide variety of inflammatory and infectious diseases and tumor progression ^3, 4^. Induction of necroptosis upon tumor necrosis factor (TNF) receptor 1 (TNFR1) activation relies on recruitment of the pro-survival TNFR1 complex I consisting of receptor-interacting protein kinase 1 (RIPK1), TNFR1-associated death domain protein (TRADD), TNFR-associated factor 2 (TRAF2), cellular inhibitor of apoptosis 1/2 (cIAP1/2) and the linear ubiquitin chain assembly complex (LUBAC) ^3^. cIAP1/2- and LUBAC-mediated K63 and M1 poly-Ub of complex I components, including RIPK1, initiate transforming growth factor β (TGFβ)-activated kinase 1 (TAK1) complex- and inhibitor of κB kinase (IKK) complex-dependent activation of pro-inflammatory mitogen-activated protein kinase (MAPK) and nuclear factor-κB (NF-κB) signaling ^5–9^. Following complex I destabilization through cIAP1/2 depletion and upon caspase-8 inhibition or depletion, the formation of a cytosolic necrosome complex is triggered ^3^. The necrosome is an amyloid-like structure composed of RIPK1 and RIPK3, interacting via their RIP homotypic interaction motif (RHIM) domains, leading to their activation by cross- and auto-phosphorylation ^10–13^. Eventually, phosphorylated RIPK3 recruits and phosphorylates MLKL at T357 and S358, resulting in the exposure of the MLKL 4-helical bundle (4HB) domain, which induces MLKL oligomerization and subsequent translocation to the plasma membrane ^14–20^. Activated MLKL accumulates in hotspots to control plasma membrane disruption and ion channel fluxes, ultimately leading to necroptotic cell death ^17, 18, 21–23^.

Necroptosis is controlled by checkpoints that include compartimentalization and species-specific post-translational modifications of RIPK1, RIPK3 and MLKL, like phosphorylation and ubiquitination ^24–26^. Ubiquitination regulates protein function, stability and localization by covalent modification of substrates with poly-Ub chains, linked via K6, K11, K27, K29, K33, K48, K63, or M1-linked (linear) Ub ^27–29^. M1 poly-Ub is mediated by the LUBAC E3 ligase, consisting of the catalytic subunit HOIP (RNF31), HOIL-1 (RBCK1) and Sharpin ^30–33^. HOIP belongs to the really interesting new gene (RING)/in-between RING (IBR)/RING (RBR)-type family of E3 ligases and specifically generates M1 poly-Ub chains through a RING-HECT hybrid reaction by transferring the activated Ub to its active site C885 and subsequently conjugating it to the acceptor Ub located in the HOIP linear ubiquitin chain determining domain (LDD) ^34–36^. The catalytic activity of HOIP can be blocked by HOIPIN-8, a small-molecule inhibitor that covalently binds to the HOIP catalytic C885 and interferes with several residues in the LDD of HOIP, thereby inhibiting the RING-HECT hybrid reaction ^37, 38^.

LUBAC generates pro-inflammatory and -survival signaling checkpoints upon activation of immune receptors, such as TNFR1, CD40, IL-1R, nucleotide oligomerization domain (NOD)-like receptors (NLRs) and toll-like receptors (TLRs) via M1 poly-Ub of receptors, receptor-associated proteins or existing K63 poly-Ub chains ^5, 6, 39^. M1 poly-Ub chains act as a scaffold to recruit signaling complexes, such as the TAK1 complex and the IKK complex to activate downstream MAPK and NF-κB signaling ^6, 8, 9, 39, 40^. Accordingly, LUBAC deficiency in mice, or in patients with genetic mutations in *HOIP*, *HOIL-1* or *Sharpin* results in inflammatory disorders linked to deregulated NF-κB activity leading to decreased gene-activatory signaling ^30–32, 41–45^. Similarly, genetic mouse models lacking functional LUBAC are sensitized towards TNFR1- and RIPK1-dependent cell death ^30, 31, 46–50^ and HOIPINs mimic LUBAC loss-of-function mutations by inhibiting cytokine-induced canonical NF-κB signaling and by regulating TNFα-induced cell viability ^37, 51, 52^.

Apart from anti-necroptotic ubiquitination, also the removal of oligomerized MLKL from the plasma membrane via endosomal sorting complex required for transport (ESCRT)-mediated plasma membrane shedding and exocytic release of MLKL or via endocytosis can counteract necroptotic cell death ^22, 53^. The homologous Flotillin-1 and -2 (FLOT1/2) proteins reside in lipid rafts, highly ordered membrane structures which are enriched in sphingolipids, cholesterol and gangliosides in the plasma membrane, the endoplasmic reticulum (ER), the Golgi apparatus and endocytic compartments ^54–57^. FLOT1/2 associate with cholesterol-rich membrane domains in the inner leaflet of cell membranes via the N-terminal stomatin, prohibitin, flotillin, HflK/C (SPFH) domain and oligomerize to heterotetramers through the C-terminal flotillin domain to create FLOT platforms ^58, 59^. FLOTs mediate the regulation of cytoskeleton-membrane interactions and cell-cell contacts, clathrin-independent endocytosis and the establishment of protein complexes at the plasma membrane ^60–68^. Interestingly, FLOTs have recently been implicated in the protection from necroptotic cell death through endocytic removal of activated MLKL from the plasma membrane ^53^.

Despite recently acquired insights into the checkpoints that control the pro-necroptotic functions of activated MLKL, it is still vastly unexplored how necroptosis is regulated downstream of MLKL activation and oligomerization. In this study, we identify LUBAC-mediated M1 poly-Ub as novel major and species-specific checkpoint in necroptotic cell death in human cells. Suppression of LUBAC and M1 poly-Ub blocks TNFα-induced necroptosis without affecting necroptotic phosphorylation of RIPK1, RIPK3 or MLKL, necrosome formation and MLKL oligomerization. Loss of M1 poly-Ub suppresses MLKL membrane hotspot accumulation, as well as MLKL-dependent release of pro-inflammatory signaling molecules, as determined by global Massive Analyses of cDNA Ends sequencing (MACE-seq)-based transcriptome analysis upon induction of necroptosis. We further identify FLOT1/2 as putative M1 poly-Ub targets during necroptosis and demonstrate that loss of FLOT1/2 additionally decreases necroptotic cell death associated with LUBAC inhibition. Finally, we confirm that loss of LUBAC activity prevents necroptosis in primary human pancreatic organoids (hPOs). Taken together, we identify a novel, species-specific role for LUBAC and M1 poly-Ub in regulating membrane accumulation of activated MLKL and promoting necroptosis. In addition, by modelling necroptotic cell death and LUBAC function in primary human organoids, we provide a novel experimental platform to study programmed cell death in intact human multicellular systems.

## Results

### LUBAC-mediated M1 poly-Ub is required for TNFα-induced necroptosis in human cells

Necroptotic cell death is tightly regulated and controlled by several molecular checkpoints, including receptor stimulation, phosphorylation of RIPK1, RIPK3 and MLKL as well as compartimentalization of phosphorylated MLKL ^22, 24, 53^. To investigate how LUBAC and M1 poly-Ub regulate these necroptotic checkpoints, cell death was induced by TNFα (T), the Smac mimetic BV6 (B) and the pan-caspase inhibitor zVAD.fmk (Z). As anticipated, prominent necroptosis could be triggered by TBZ in human colon carcinoma HT-29 cells, as determined by high-content imaging-based quantification of propidium iodide (PI)-positive cells **(Figure 1A)**. HT-29 cells are highly sensitive towards TNFα-induced programmed cell death and are commonly used model cell lines to study necroptosis. Surprisingly, chemical inhibition of HOIP, the catalytic subunit of LUBAC, with the small molecular weight inhibitor HOIPIN-8 ^38^, rescued TBZ-induced necroptosis in a time-dependent manner **(Figure 1A)**. Since LUBAC subunit levels are differentially regulated in myeloid cells as compared to other cell types ^69, 70^, the effect of HOIP inhibition was investigated in the human monocytic cell line THP-1. TBZ-induced necroptotic cell death in THP-1 cells could also be potently reversed by HOIPIN-8 **(Figure 1B)**, confirming a broader relevance of HOIPIN-8-mediated inhibition in necroptosis. In agreement with HOIPIN-8-mediated necroptosis inhibition, chemical inhibition of LUBAC activity with the mycotoxin gliotoxin ^71^ also effectively blocked necroptotic cell death, without inducing additional cytotoxicity **(Figures S1A and S1B)**. Importantly, the effects of HOIPIN-8- and gliotoxin-based HOIP inhibition on necroptosis could be confirmed by siRNA-mediated knock-down of HOIP expression that rescued necroptosis to a similar extent as HOIPIN-8- and gliotoxin-mediated LUBAC inhibition **(Figures 1C and S1C)**, making off-target effects unlikely. Importantly, total cellular levels of M1 poly-Ub increased upon induction of necroptosis, which could be efficiently reduced by HOIPIN-8 and siRNA-mediated knock-down of HOIP in HT-29 cells **(Figures 1D and 1E)** and THP-1 cells **(Figure S1D)**. Together, these observations reveal that LUBAC-mediated M1 poly-Ub serves as a novel central checkpoint and conserved mediator of TNFα-induced necroptotic cell death.

**Figure 1.**
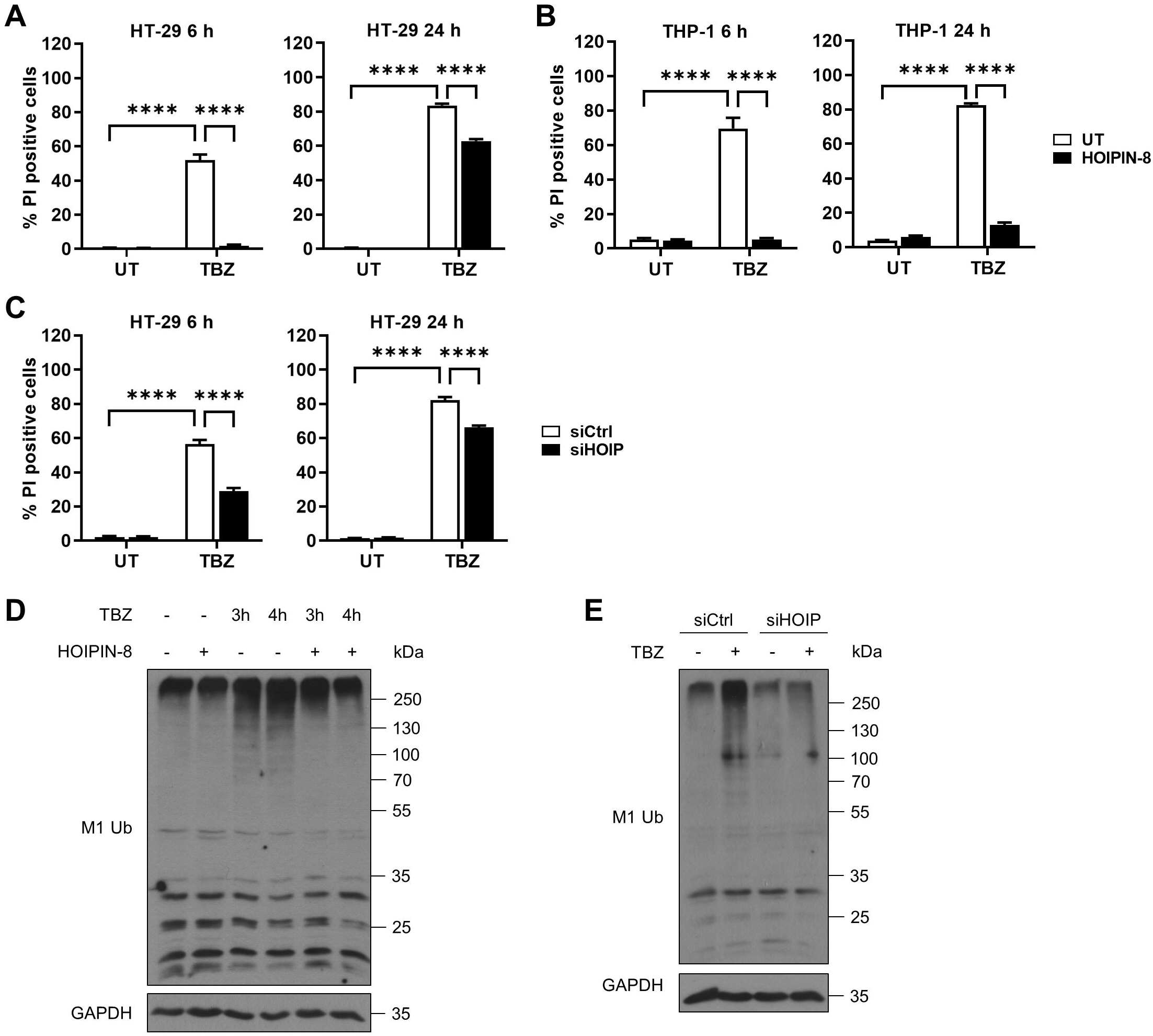
LUBAC-mediated M1 poly-Ub is required for TNFα-induced necroptosis in human cells. **A.** Quantification of cell death in untreated (UT) and HOIPIN-8 (30 µM) pre-treated HT-29 cells after treatment with TBZ (10 ng/mL TNFα, 1 µM BV6, 20 µM zVAD.fmk) for the indicated time points. Mean and SEM of at least three independent experiments are shown. \*\*\*\**P*<0.0001. **B.** Quantification of cell death in untreated (UT) and HOIPIN-8 (30 µM)-pre-treated THP-1 cells after treatment with TBZ (10 ng/mL TNFα, 1 µM BV6, 20 µM zVAD.fmk) for the indicated time points. Mean and SEM of at least three independent experiments are shown. \*\*\*\**P*<0.0001. **C.** Quantification of cell death in control (siCtrl) and HOIP knockdown (siHOIP) HT-29 cells treated with TBZ (10 ng/mL TNFα, 1 µM BV6, 20 µM zVAD.fmk) for the indicated time points. Mean and SEM of at least three independent experiments are shown. \*\*\*\**P*<0.0001. **D.** Western blot analysis of total M1 poly-Ub levels in control or HOIPIN-8 (30 µM)-pre-treated HT-29 cells upon treatment with TBZ (10 ng/mL TNFα, 1 µM BV6, 20 µM zVAD.fmk) for the indicated time points. GAPDH was used as loading control. Representative blots of at least two independent experiments are shown. **E.** Western blot analysis of total M1 poly-Ub levels in control (siCtrl) and HOIP knockdown (siHOIP) HT-29 treated with TBZ (10 ng/mL TNFα, 1 µM BV6, 20 µM zVAD.fmk) for 4 h. GAPDH was used as loading control. Representative blots of at least two independent experiments are shown. See also Figure S1.

Up till now, the relevance of M1 poly-Ub generated by LUBAC in TNFα-mediated cell fate signaling has been prominently demonstrated in genetic mouse models deficient in *HOIP*, *HOIL-1* or *Sharpin*. Loss of either HOIP, HOIL-1 or Sharpin in mice interferes with TNFR1-mediated signaling, leading to impaired NF-κB activation and increased sensitivity towards TNFR1-mediated programmed cell death ^30, 46–50, 72, 73^. Therefore, the consequences of LUBAC inhibition on TNFα-induced necroptosis were evaluated in mouse embryonic fibroblasts (MEFs) and L-929 mouse lung fibroblasts. HOIPIN-8 acts by interfering with the active site residue C885 as well as residues in the HOIP LDD ^37^, which are conserved in humans and mice **(Figure S1E)**. LUBAC inhibition efficiently abrogated TNFα-induced NF-κB activation and stabilized phosphorylated IκBα in both HT-29 and MEFs **(Figures S1F and S1G)**, consistent with the described pro-inflammatory roles of LUBAC and M1 poly-Ub and in agreement with the abovementioned findings. However, to our surprise, HOIPIN-8-mediated LUBAC inhibition did not rescue TBZ-induced cell death in MEFs **(Figure S1H)** and even enhanced TBZ-induced cell death in L-929 cells, without inducing cytotoxicity, **(Figure S1I)**, which could be effectively blocked by inhibition of RIPK3 with GSK’872 **(Figure S1I)**. This is in accordance with previous findings demonstrating that RNAi-mediated silencing of HOIP or HOIL-1 sensitizes L-929 cells to necroptotic cell death ^73^. Of note, the cellular levels of M1 poly-Ub remained relatively stable upon necroptosis induction in MEFs **(Figure S1J)**. These observations, confirmed by a detailed analysis of cell death regulation in genetic mouse models of LUBAC deficiency ^30, 46, 47, 72^, suggest that LUBAC acts as species-specific switch in the control of necroptosis.

### Loss of LUBAC-catalyzed M1 poly-Ub does not affect necroptotic signaling and necrosome formation

To mechanistically understand how LUBAC controls necroptosis, the role of M1 poly-Ub on necroptotic checkpoints was analyzed in further detail. Importantly, necroptosis induction did not affect the expression levels of the LUBAC subunits HOIP, HOIL-1 and Sharpin **(Figures 2A and S2A)**, suggesting that LUBAC activity is responsible for the increased levels of M1 poly-Ub during necroptosis, instead of alterations in LUBAC abundance. In addition, HOIPIN-8-mediated inhibition of LUBAC did not have any influence on the expression patterns of individual LUBAC subunits, excluding the possibility that selective loss of subunits might cause the observed HOIPIN-8-induced effects on M1 poly-Ub during necroptosis **(Figures 2A and S2A)**.

**Figure 2.**
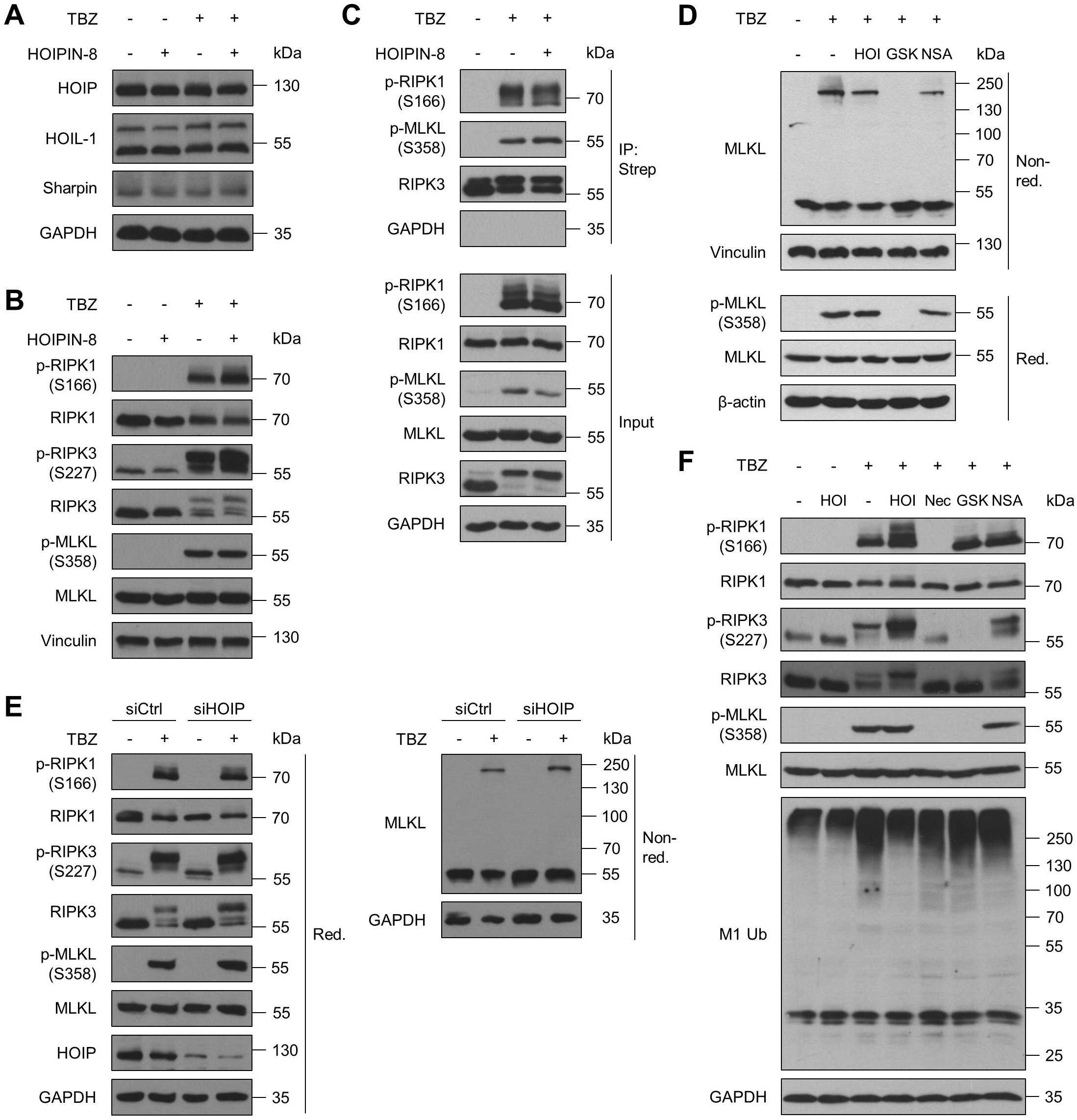
Loss of LUBAC-catalyzed M1 poly-Ub does not affect necroptotic signaling and necrosome formation. **A.** Western blot analysis of LUBAC subunit (HOIP, HOIL-1 and Sharpin) expression levels in control *versus* HOIPIN-8 (30 µM)-pre-treated HT-29 cells upon treatment with TBZ (10 ng/mL TNFα, 1 µM BV6, 20 µM zVAD.fmk) for 4 h. GAPDH was used as loading control. Representative blots of at least two independent experiments are shown. **B.** Western blot analysis of phosphorylated and total expression levels of RIPK1, RIPK3 and MLKL in control *versus* HOIPIN-8 (30 µM)-pre-treated HT-29 cells upon treatment with TBZ (10 ng/mL TNFα, 1 µM BV6, 20 µM zVAD.fmk) for 4 h. Vinculin was used as loading control. Representative blots of at least two independent experiments are shown. **C.** Immunoprecipitation of Strep-RIPK3 from control and HOIPIN-8 (30 µM)-pre-treated RIPK3 KO HT-29 cells re-expressing PAM-mutated doxycycline (Dox)-inducible Strep-tagged RIPK3 wild-type (WT) was performed in order to analyze necrosome formation after treatment with TBZ (10 ng/mL TNFα, 1 µM BV6, 20 µM zVAD.fmk) for 3 h. Cells were incubated overnight with 1 μg/mL Dox prior to treatment to induce expression of Strep-RIPK3. GAPDH was used as loading control. Representative blots of at least two independent experiments are shown. **D.** Western blot analysis of MLKL oligomerization upon treatment with TBZ (10 ng/mL TNFα, 1 µM BV6, 20 µM zVAD.fmk) for 4 h by non-reducing Western blotting of control, HOIPIN-8 (30 µM), GSK’872 (20 µM) or NSA (10 µM)-pre-treated HT-29 cells. Vinculin or β-actin were used as loading controls. Representative blots of at least two independent experiments are shown. **E.** Western blot analysis of expression levels of phosphorylated and total RIPK1, RIPK3 and MLKL and MLKL oligomerization (by non-reducing Western blotting) upon 4 h TBZ (10 ng/mL TNFα, 1 µM BV6, 20 µM zVAD.fmk) treatment of control (siCtrl) and HOIP knockdown (siHOIP) HT-29 cells. GAPDH was used as loading control. Representative blots of at least two independent experiments are shown. **F.** Western blot analysis of expression levels of phosphorylated and total RIPK1, RIPK3, MLKL and M1 Ub upon 4 h TBZ (10 ng/mL TNFα, 1 µM BV6, 20 µM zVAD.fmk) treatment of HT-29 cells pre-treated with HOIPIN-8 (30 µM) or Nec-1s (30 µM), GSK‘872 (20 µM) and NSA (10 µM). GAPDH was used as loading control. Representative blots of at least two independent experiments are shown. See also Figure S2.

Surprisingly, and in contrast to the prominent HOIPIN-8-mediated rescue of necroptosis, HOIPIN-8 did not attenuate TBZ-induced phosphorylation of RIPK1 at S166, RIPK3 at S227 and MLKL at S358, major checkpoints in necroptotic cell death signaling **(Figure 2B and Figures S2B and S2C)**. In contrast, LUBAC inhibition induced a slight accumulation of phosphorylated RIPK1 S166, RIPK3 S227 and MLKL S358 **(Figures 2B and Figures S2B and S2C)**, suggesting alternative effects of LUBAC on the core necroptotic cell death machinery.

A central hallmark of necroptosis consists in the formation of the necrosome, a specialized amyloid-like signaling platform, composed of activated RIPK1 and RIPK3, that eventually phosphorylates MLKL to drive necroptotic cell death ^10, 11, 14, 17^. Importantly, inhibition of LUBAC did not interfere with necrosome formation, as determined by co-immunoprecipitation of phosphorylated RIPK1 S166 and MLKL S358 with RIPK3 from untreated and HOIPIN-8-treated necroptotic cells **(Figure 2C)**.

Prior to membrane accumulation, activated MLKL oligomerizes into larger structures that are required for efficient necroptosis progression ^17–20^. Upon induction of necroptosis, prominent MLKL oligomerization could be observed that, in contrast to GSK’872-mediated RIPK3 inhibition, could not be blocked by HOIPIN-8 **(Figure 2D)**. Furthermore, inhibiting MLKL with necrosulfonamide (NSA) did neither affect MLKL oligomerization nor phosphorylation **(Figure 2D)**, as NSA only prevents membrane accumulation of activated MLKL ^17^. Similarly, siRNA-mediated loss of HOIP also did not have any effect on necroptosis-induced RIPK1 S166, RIPK3 S227 and MLKL S358 phosphorylation and did not attenuate MLKL oligomerization **(Figure 2E)**. In contrast to HOIPIN-8-mediated inactivation of LUBAC, inhibition of RIPK1 with necrostatin-1 (Nec-1s) or RIPK3 with GSK’872 influences necroptosis-induced phosphorylation of RIPK1 S166, RIPK3 S227 and MLKL S358, respectively, while blocking MLKL with NSA did not affect phosphorylation levels **(Figure 2F)**. Of note, in contrast to HOIPIN-8, Nec-1s, GSK’872 or NSA, as well as deletion of RIPK3 or MLKL, did not influence total M1 poly-Ub levels upon induction of necroptosis, suggesting that regulation of M1 poly-Ub occurs in parallel with necroptosis initiation and progression **(Figures 2F and S2C)**.

### LUBAC-mediated M1 poly-Ub regulates necroptosis-induced and MLKL-dependent production of inflammatory signaling molecules

Upon the induction of necroptosis, activated MLKL triggers the transcriptional upregulation and secretion of a variety of inflammatory signaling molecules, including cytokines and chemokines, like CXCL1 and CXCL10 ^1, 22^. Consistently with these findings, TBZ-triggered necroptosis strongly induced expression of CXCL1 and CXCL10 mRNA **(Figure 3A)** in an MLKL-dependent manner, since inhibition of MLKL with NSA and CRISPR/Cas9-mediated knock-out (KO) of MLKL expression largely prevented necroptosis-induced upregulation of CXCL1 and CXCL10 mRNA (**Figures 3A and S3A)**. Interestingly, LUBAC inhibition in necroptotic cells reduced CXCL1 and CXCL10 mRNA expression **(Figure 3A)** and similar findings were observed upon loss of HOIP expression **(Figure 3B)**. In agreement with the HOIP-dependent alterations in CXCL1 and CXCL10 mRNA expression, secretion of CXCL1 during necroptosis was also diminished in the presence of HOIPIN-8 or upon loss of HOIP expression **(Figures 3C and 3D)**. To further characterize the LUBAC-dependent regulation of the transcriptional changes associated with necroptosis, transcriptome profiling using unbiased MACE-seq was perfomed in necroptotic cells upon inhibition of LUBAC activity. Necroptosis prominently triggered the transcriptional upregulation of a highly pro-inflammatory gene expression profile **(Figure 3E)**, in agreement with previous observations ^1^. Importantly, loss of LUBAC activity potently reversed the necroptotic inflammatory profile and strongly attenuated the expression of necroptosis-upregulated cytokines and chemokines, such as TNFα, CSF1, CXCL2 and CCL20, as well as the adhesion molecule ICAM1 **(Figure 3E, Figures S3B and S3C)**. Of note, necroptosis-induced expression of TNFα and ICAM1 could be largely prevented by inhibiton of MLKL with NSA **(Figures S3B and S3C)**, confirming that MLKL is required for necroptosis-induced upregulation of regulatory and inflammatory gene expression signatures.

**Figure 3.**
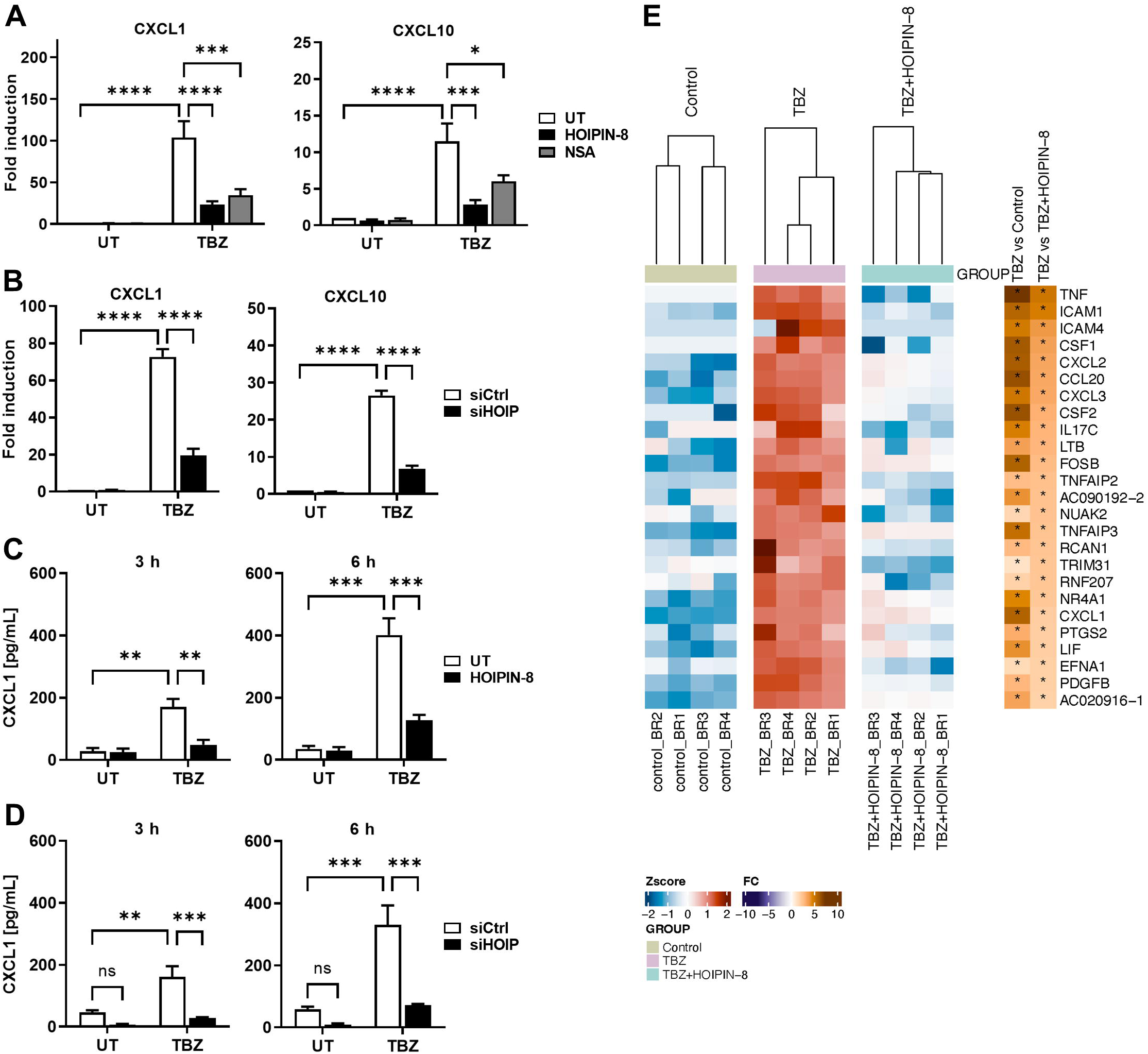
LUBAC-mediated M1 poly-Ub regulates necroptosis-induced and MLKL-dependent production of inflammatory signaling molecules. **A.** mRNA expression levels of CXCL1 and CXCL10 of control (UT), HOIPIN-8 (30 µM) or NSA (10 µM)-pre-treated HT-29 cells upon treatment with TBZ (10 ng/mL TNFα, 1 µM BV6, 20 µM zVAD.fmk) for 3 h. Gene expression was normalized against 18S and RPII mRNA expression and is presented as x-fold mRNA expression compared to the untreated (UT) control. Mean and SEM of at least three independent experiments are shown. \**P*<0.05; \*\*\**P*<0.001; \*\*\*\**P*<0.0001. **B.** mRNA expression levels of CXCL1 and CXCL10 of control (siCtrl) or HOIP knockdown (siHOIP) HT-29 cells treated with TBZ (10 ng/mL TNFα, 1 µM BV6, 20 µM zVAD.fmk) for 4 h. Gene expression was normalized against 18S and RPII mRNA expression and is presented as x-fold mRNA expression compared to the untreated (UT) control cells. Mean and SEM of at least three independent experiments are shown. \*\*\*\**P*<0.0001. **C.** ELISA-based quantification of CXCL1 secretion in supernatants of control (UT) or HOIPIN-8 (30 µM)-pre-treated HT-29 cells after treatment with TBZ (10 ng/mL TNFα, 1 µM BV6, 20 µM zVAD.fmk) for the indicated time points. Mean and SEM of at least three independent experiments are shown. \*\**P*<0.01; \*\*\**P*<0.001. **D.** ELISA-based quantification of CXCL1 secretion in supernatants of control (siCtrl) or HOIP knockdown (siHOIP) HT-29 cells treated with TBZ (10 ng/mL TNFα, 1 µM BV6, 20 µM zVAD.fmk) for the indicated time points. Mean and SEM of at least three independent experiments are shown. \*\**P*<0.01; \*\*\**P*<0.001. **E.** MACE-seq-determined heatmap showing the row-wise scaled intensity (left panel) of the top-25 upregulated genes in HT-29 cells treated with TBZ (10 ng/mL TNFα, 1 µM BV6, 20 µM zVAD.fmk) for 3 h in comparison to HOIPIN-8 (30 µM)-pre-treated and TBZ-treated HT-29 cells. Genes are sorted according to their log2 fold change values (right panel). \**P*<0.05. See also Figure S3.

Together, these data suggest that M1 poly-Ub is required for the activated MLKL-dependent secretion of signaling molecules, cytokines and chemokines, such as CXCL1 and CXCL10, upon induction of necroptosis.

### LUBAC and M1 poly-Ub segregate the subcellular distribution of activated MLKL

During the effector phase of necroptosis, activated MLKL oligomers grow to form cytoplasmic clusters which translocate to the cell membrane to accumulate into hotspots, preferentially at intercellular junctions ^21^. To investigate whether LUBAC-mediated M1 poly-Ub might control membrane translocation of activated MLKL, a non-ionic detergent- and temperature-dependent fractionation of micelle-poor (aqueous) and micelle-rich (detergent-enriched) fractions was applied to isolate hydrophobic, membrane-associated proteins from aqueous solutions. In necroptotic cells, phosphorylated MLKL accumulated in detergent-enriched fractions, which could be efficiently blocked by NSA **(Figure 4A)**, suggesting extensive membrane accumulation that is consistent with previous findings ^17^. Surprisingly, LUBAC inhibition prevented accumulation of phosphorylated MLKL in detergent-enriched fractions, leading instead to an enrichment of phosphorylated MLKL in the aqueous fraction **(Figure 4A)**. These data suggest that LUBAC-mediated M1 poly-Ub is a prerequisite for the translocation of activated MLKL from the cytosol to membrane compartments during necroptosis induction. Of note, necroptosis also triggered the accumulation of M1 poly-Ub chains in both aqueous and detergent-enriched fractions, that could be counteracted by LUBAC inhibition **(Figure S4A)**.

**Figure 4.**
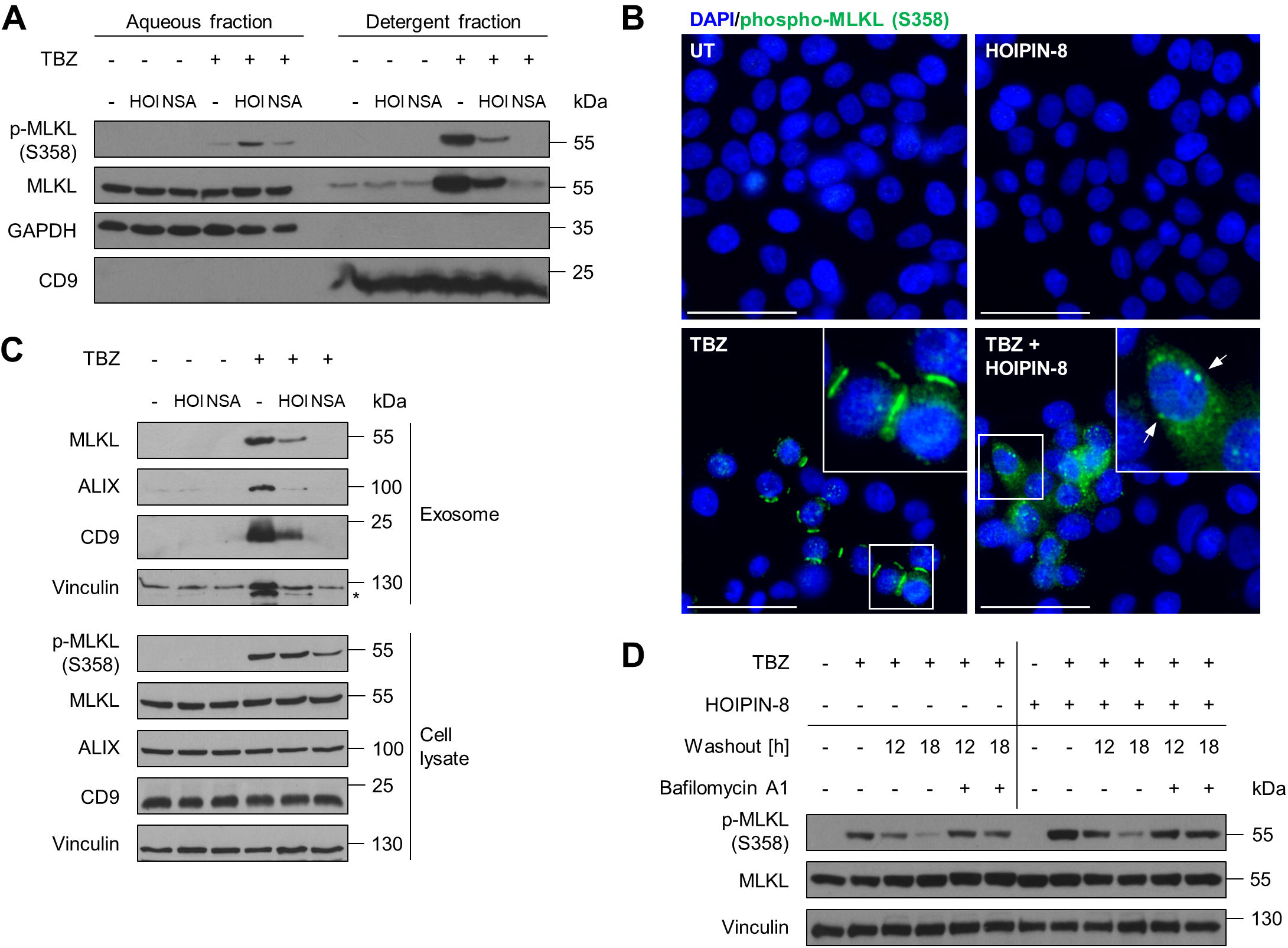
LUBAC and M1 poly-Ub segregate the subcellular distribution of activated MLKL. **A.** Fractionation of phosphorylated and total MLKL in the micelle-poor (aqueous) and micelle-rich (detergent) fractions upon phase separation using Triton X-114 lysis buffer in untreated (UT) and HOIPIN-8 (30 µM) pre-treated HT-29 cells upon treatment with TBZ (10 ng/mL TNFα, 1 µM BV6, 20 µM zVAD.fmk) for 4 h. GAPDH was used as loading control for soluble proteins and CD9 as loading control for membrane proteins. Representative blots of at least two independent experiments are shown. **B.** Representative fluorescence microscopy demonstrating the cellular distribution of phosphorylated MLKL (green) in untreated (UT) and HOIPIN-8 (30 µM) pre-treated HT-29 cells and after treatment with TBZ (10 ng/mL TNFα, 1 µM BV6, 20 µM zVAD.fmk) for 3 h. Nuclei were stained with DAPI (blue). Arrows indicate cytoplasmic clusters of phosphorylated MLKL. Representative images of at least two independent experiments are shown. Scale bar 50 µm. **C.** Western blot analysis of TBZ-induced release of MLKL in exosomes, isolated from the supernatants of control, HOIPIN-8 (30 µM) or NSA (10 µM)-pre-treated HT-29 cells after treatment with TBZ (10 ng/mL TNFα, 1 µM BV6, 20 µM zVAD.fmk) for 3 h. Exosome isolation was confirmed with antibodies against ALIX, CD9 and Vinculin. Asterisk marks unspecific band. Expression of phosphorylated and total MLKL was determined by Western blotting of cell lysates from the same experiment. Vinculin was used as loading control. Representative blots of at least two independent experiments are shown. **D.** Western blot analysis of lysosomal degradation of phosphorylated MLKL upon TBZ (10 ng/mL TNFα, 0.1 µM BV6, 20 µM zVAD.fmk) treatment of control, HOIPIN-8 (30 µM) or NSA (10 µM)-pre-treated HT-29 cells. HT-29 cells were treated with TBZ for 3 h and either harvested or the medium was exchanged with fresh medium with or without BafA1 (100 nM) after a washing step with PBS. Washed-out cells were harvested 12 h or 18 h after medium exchange. Vinculin was used as loading control. Representative blots of at least two independent experiments are shown. See also Figure S4.

To further understand how LUBAC and M1 poly-Ub regulate the membrane localization of phosphorylated MLKL during necroptosis, MLKL distribution was assessed by immunofluorescence. Upon necroptosis induction, S358-phosphorylated MLKL localized to membranes in large hotspots at intercellular junctions **(Figures 4B and S4B)**, corresponding with previous findings ^21^. As anticipated, MLKL inhibition with NSA blocked membrane accumulation of activated MLKL in necroptotic cells **(Figure S4B)**. In accordance with the altered cellular distribution of MLKL in fractionation experiments, HOIPIN-8 completely abrogated the formation of phosphorylated MLKL hotspots at the plasma membrane in necroptotic cells, leading to the appearance of diffuse cytoplasmic punctate clusters of S358 phosphorylated MLKL **(Figure 4B)**.

The cellular fate of phosphorylated MLKL determines the functional outcome of necroptotic signaling and the regulation of the availability of necroptosis-prone phosphorylated MLKL is tightly controlled. At one end, activated MLKL is internalized through endocytosis and degraded through the lysosomal pathway ^53^. At the other end, the levels of activated MLKL are controlled by exocytosis into extracellular vesicles ^22, 53, 74, 75^. TBZ-induced necroptosis strongly triggered exocytosis leading to the formation of ALIX-, CD9- and vinculin-positive extracellular exosomes that contained high levels of MLKL **(Figure 4C)**. HOIPIN-8 strongly reduced the necroptosis-associated formation of extracellular vesicles and decreased the accumulation of MLKL in exosomes **(Figure 4C)**. As expected, MLKL inhibition with NSA completely blocked TBZ-induced exosome formation and release of MLKL **(Figure 4C)**. In line with these observations, HOIPIN-8-mediated inhibition of LUBAC prevented lysosomal degradation of phosphorylated MLKL upon necroptosis induction, as determined by washout of the necroptotic stimulus in the absence or presence of the lysosomal inhibitor bafilomycin A1 (BafA1) **(Figure 4D)**. Intriguingly, LUBAC inhibition in necroptotic cells increased levels of phosphorylated MLKL S358 **(Figure 4D),** corresponding with previous data **(Figure S2B)**. Taken together, these data demonstrate that LUBAC-mediated M1 poly-Ub regulates necroptosis downstream of MLKL phosphorylation and oligomerization by controlling the availability and translocation of activated MLKL to membranes and MLKL hotspot formation at the plasma membrane.

### FLOT1/2 license necroptosis in a M1 poly-Ub-dependent manner

Recent studies revealed that FLOT1/2 colocalize with MLKL during necroptosis and act as important mediators of MLKL endocytosis ^53, 75^. To understand the potential regulatory link between M1 poly-Ub, the cellular distribution of activated MLKL and FLOT1/2, enrichment of M1 poly-Ub chains, substrates and associated proteins was performed from necroptotic cells. Interestingly, high levels of FLOT1/2 could be enriched by capture of M1 poly-Ub via ubiquitin binding in ABIN and NEMO (UBAN)-based pulldowns from necroptotic cells **(Figures 5A and S5A)**. Of note, HOIPIN-8-mediated LUBAC inhibition completely prevented M1 poly-Ub-associated enrichment of FLOT1/2, without affecting total FLOT1/2 cellular expression levels **(Figures 5A and S5A)**. Furthermore, loss of FLOT1 or -2 individually, or combined loss of FLOT1/2, decreased necroptotic cell death and further potentiated the repressive effects of LUBAC inhibition on necroptosis **(Figures 5B and S5B)**. Loss of FLOT1/2 did not affect necroptosis-induced increases in M1 poly-Ub or HOIPIN-8-mediated decreases in necroptosis-triggered M1 poly-Ub **(Figure S5C)**. In addition, combined loss of FLOT1/2 did not interfere with necrosome formation **(Figure 5C)** and did not affect necroptosis-induced MLKL oligomerization in the absence or presence of HOIPIN-8 **(Figure 5D)**. Intruigingly, enrichment of FLOT1 in UBAN-based pulldowns could be prevented by inhibition of RIPK1 with Nec-1s, RIPK3 with GSK’872 and MLKL with NSA, indicating that activation and membrane translocation of MLKL is a prerequisite for association of FLOT1 with M1 poly-Ub **(Figure 5E)**.

**Figure 5.**
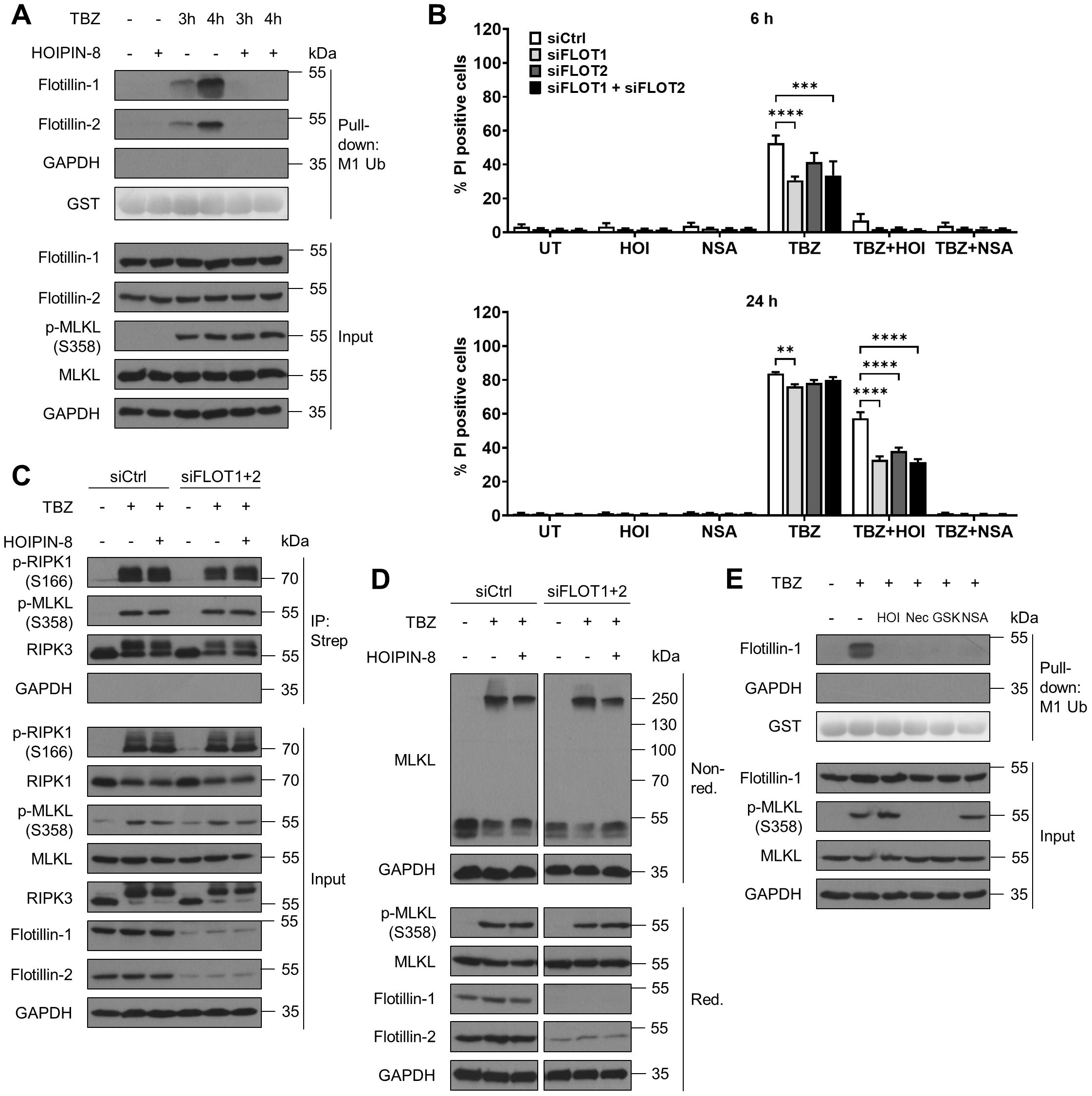
FLOT1/2 license necroptosis in a M1 poly-Ub-dependent manner. **A.** UBAN-mediated pulldown of M1 ubiquitinated proteins in control or HOIPIN-8 (30 µM)-pre-treated HT-29 cells treated with TBZ (10 ng/mL TNFα, 1 µM BV6, 20 µM zVAD.fmk) for the indicated time points. GST and GAPDH were used as loading controls. Representative blots of at least two independent experiments are shown. **B.** Quantification of cell death in untreated (UT), HOIPIN-8 (30 µM) or NSA (10 µM)-pre-treated control (siCtrl), FLOT1 (siFLOT1), FLOT2 (siFLOT2) or combined FLOT1 and FLOT2 (siFLOT1+siFLOT2) knockdown HT-29 cells after treatment with TBZ (10 ng/mL TNFα, 1 µM BV6, 20 µM zVAD.fmk) for the indicated time points. Mean and SEM of at least three independent experiments are shown. \*\**P*<0.001; \*\*\**P*<0.001; \*\*\*\**P*<0.0001. **C.** Characterization of necrosome formation by immunoprecipitation of Strep-RIPK3 from control and HOIPIN-8 (30 µM)-pre-treated control (siCtrl) and combined FLOT1 and FLOT2 knockdown (siFLOT1+2) RIPK3 KO HT-29 cells re-expressing PAM-mutated Dox-inducible Strep-tagged RIPK3 WT after treatment with TBZ (10 ng/mL TNFα, 1 µM BV6, 20 µM zVAD.fmk) for 3 h. Cell lines were incubated overnight with 1 μg/mL Dox prior to treatment to induce expression of Strep-RIPK3. GAPDH was used as loading control. Representative blots of at least two independent experiments are shown. **D.** Western blot analysis of MLKL oligomerization after treatment with TBZ (10 ng/mL TNFα, 1 µM BV6, 20 µM zVAD.fmk) for 4 h by non-reducing Western blotting of control (siCtrl) and combined FLOT1 and FLOT2 knockdown (siFLOT1+2) HT-29 cells. GAPDH was used as loading control. Representative blots of at least two independent experiments are shown. **E.** UBAN-mediated pulldown of M1 ubiquitinated proteins in control, HOIPIN-8 (30 µM), Nec-1s (30 µM), GSK‘872 (20 µM) or NSA (10 µM)-pre-treated HT-29 cells treated with TBZ (10 ng/mL TNFα, 1 µM BV6, 20 µM zVAD.fmk) for 4 h. GST and GAPDH were used as loading controls. Representative blots of at least two independent experiments are shown. See also Figure S5.

### LUBAC-mediated M1 poly-Ub regulates necroptotic cell death in primary human pancreatic organoids

Our data reveal that LUBAC and M1 poly-Ub act as a novel, species-specific checkpoint that control the membrane hotspot accumulation of activated MLKL and thereby regulate necroptotic cell death in human, but not in murine cell lines. To further confirm this concept and to exclude artefacts in necroptotic signaling related to the use of human cancer cell lines, LUBAC- and M1 poly-Ub-mediated regulation of necroptosis was studied in primary human organoids. Organoids are stem cell-derived three-dimensional cellular clusters consisting of organ-specific cell types that self-organize and exhibit similar functionality and complexity as their tissue of origin ^76, 77^. Organoids represent highly physiological models as compared to two-dimensional monolayer cell culture models ^78^. To model necroptosis, adult stem cell-derived primary human pancreatic organoids (hPOs) recapitulate the phenotype of the original pancreatic ductal epithelia and retain the pancreatic morphology and the distribution of ductal, acinar, mesenchymal and endothelial cells in a hollow spherical polarized cell monolayer enclosing a central lumen ^79–82^. hPOs were maintained in matrix for 3 to 5 days to allow the formation of organoid spheres composed of a cellular monolayer surrounding a liquid-filled lumen, that expanded over time **(Figures 6A and S6A)**, in line with previous findings ^83^, after which necroptosis was induced. For this, matrix-embedded primary hPOs were incubated with BV6 (B) and the clinically-applicable pan-caspase inhibitor IDN-6556 (Emricasan; E), followed by time-lapse imaging and subsequent fluorescein diacetate (FDA)- and PI-based live/dead imaging. BV6 combined with Emricasan induced a prominent loss of luminal hPO morphology, characterized by organoid collapse and compaction, with individual cells migrating away from the collapsed organoids **(Figure 6A, Movie 1)**. Live/dead staining of BE-incubated organoids revealed high levels of PI-positive cells and low levels of FDA-positive cells, confirming prominent hPO cell death **(Figure 6B)**. Similar organoid collapse and loss of luminal morphology was observed in hPOs incubated with the Smac mimetic Birinapant (Bi) and Emricasan **(Figure S6A, Movie 2)** and upon incubation with BV6 and zVAD.fmk (BZ) **(Figure S7)**. Importantly, HOIPIN-8-mediated inhibition of LUBAC delayed BE-induced organoid collapse, prevented the loss of luminal hPO morphology and extended hPO viability up to 24 h **(Figures 6A and 6B, Movie 1)**, although hPOs appeared shrunken and smaller in size **(Figure 6A)**. HOIPIN-8 also blocked organoid collapse and cell death induced by BiE and BZ **(Figures S6A and S6B, Figure S7, Movie 2)**. Of note, organoid collapse and compaction, loss of hPO morphology and cell death triggered by BiE could be blocked by NSA, indicating the appearance of MLKL-dependent necroptotic cell death **(Figure S6, Movie 2)**.

**Figure 6.**
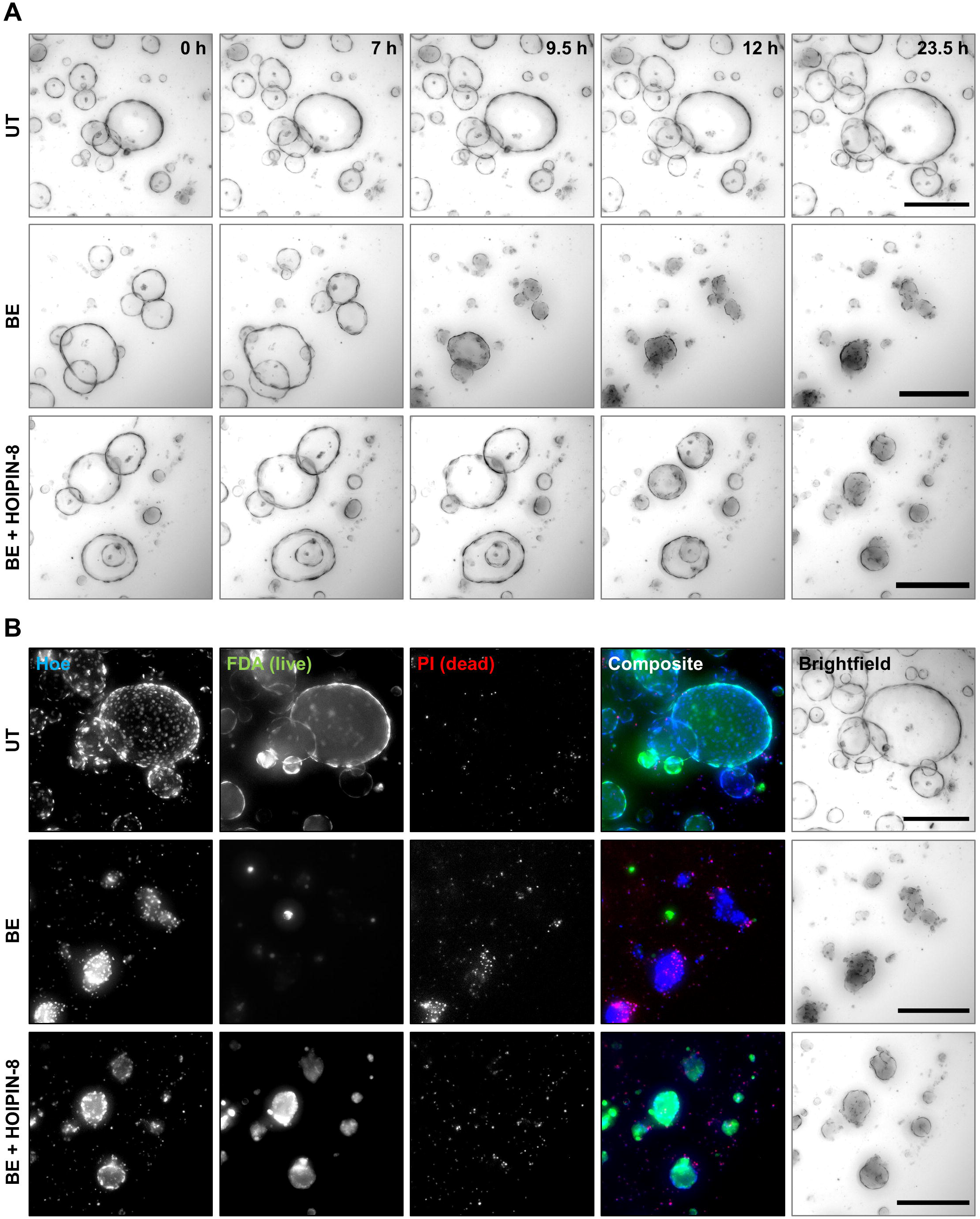
LUBAC-mediated M1 poly-Ub regulates necroptotic cell death in primary hPOs. **A.** Representative images of time-lapse videos of untreated (UT), BV6 (1 µM) and Emricasan (10 µM) (BE)-treated and HOIPIN-8 (30 µM)-pre-treated and BE (BE+HOIPIN-8)-treated primary hPOs for a total period of 23.5 h. Scalebars: 250 µm. **B.** *Idem* as A., but treated primary hPOs were stained with Hoechst33342 (Hoe; blue), FDA (live; green) and PI (dead; red) after 24 h treatment with BE or BE+HOIPIN-8 prior to imaging. Scalebars: 250 µm. See also Figures S6 and S7 and Movies 1 and 2.

In conclusion, we identify LUBAC-mediated M1 poly-Ub as novel checkpoint that regulates necroptosis by controlling the cellular distribution of activated MLKL between membrane hotspot formation and endo- and exocytosis in cell lines of human origin and primary human organoids.

## Discussion

Necroptosis relies on the highly regulated phosphorylation of RIPK1 and RIPK3 in the context of the death-inducing necrosome complex, leading to phoshorylation of the pseudokinase MLKL, followed by MLKL oligomerization, membrane accumulation and cell death ^11–15, 17^. By doing so, necroptotic cell death is involved in a wide variety of diseases, including infection ^84–87^, cancer ^88, 89^ and ischemia-reperfusion injury ^90, 91^. TNFα-induced necroptosis, but also alternatively-induced forms of necroptosis, are controlled by checkpoints that include compartimentalization, complex formation and post-translational modification, like phosphorylation and ubiquitination, of the key necroptotic proteins RIPK1, RIPK3 and MLKL ^24–26^. M1-linked poly-Ub, catalyzed by the LUBAC E3 ligase, serves as important regulator that balances pro-and anti-survival measures by modification of RIPK1. Here, we identify a novel regulatory function of M1 poly-Ub during necroptosis in facilitating the accumulation of activated MLKL in membrane hotspots to promote necroptotic cell death in cells of human origin.

In contrast to genetic models of LUBAC deficiency in mice, in which LUBAC-mediated M1 poly-Ub represses cell death ^30, 31, 46–50, 72^, we found that pharmacological inhibition of LUBAC with either HOIPIN-8 or gliotoxin, as well as siRNA-mediated loss of LUBAC, prevents TNFα-induced necroptosis in human, but not murine, cell lines. Induction of necroptosis triggered M1 poly-Ub that was not dependent on the activity or the physical presence of RIPK1, RIPK3 and MLKL, suggesting that increases in M1 poly-Ub during necroptosis are not regulated by the core necroptotic machinery. In addition, loss of LUBAC activity reverts the cellular levels of necroptosis-induced M1 poly-Ub in human cell lines, but to a lesser degree in cell lines of murine origin. LUBAC inhibition abrogated TNFα-induced NF-κB signaling to the same extent in cell lines from both species, suggesting species-, pathway- and context-specific modes of LUBAC regulation during necroptosis. Importantly, structural studies revealed important mechanistic and functional differences between human and murine necroptotic signaling pathways ^92–95^. These confirm that the mechanisms by which MLKL is activated by RIPK3 differ substantially in humans and mice, as a result of species-specific coevolution and divergence of RIPK3 and MLKL ^92, 94^. The activation of MLKL is achieved through RIPK3-dependent phosphorylation of human MLKL at T357 and S358 and murine MLKL at S345, located in the activation loop of MLKL ^14, 15, 96, 97^. Intruigingly, human and murine MLKL are not able to complement necroptotic signaling when expressed in cells of the opposing species ^92–94^. One of the most prominent differences in how human and murine MLKL are activated relies on the transient interaction between murine RIPK3 and MLKL, which is in striking contrast to the activation of human MLKL that requires recruitment and binding to RIPK3 ^92^, indicating additional roles for T357/S358 phosphorylation in human MLKL ^92^. In addition, species-specific, yet unidentified, differences in necroptosis execution downstream of activated MLKL may also explain the distinct functions of LUBAC and M1 poly-Ub in the regulation of necroptotic cell death in mice and humans.

Our findings reveal that, apart from the core necroptosis machinery, LUBAC-dependent signaling events might become activated in parallel that involve M1 poly-Ub to eventually affect necroptosis. In line with this, loss of LUBAC activity did not alter necroptotic phosphorylation of RIPK1, RIPK3 and MLKL, necrosome formation or the oligomerization of MLKL during necroptosis induction. In addition, TNFα-induced necroptosis has been linked to the cell-autonomous upregulation and increased secretion of signaling molecules, including cytokines and chemokines ^1, 22^. Induction of this necroptotic cytokine profile involves two phases, the first of which depends on TNFα-induced NF-κB-mediated upregulation of inflammatory signaling molecules. The second phase is a RIPK1-, RIPK3- and MLKL-mediated wave, that relies on necroptotic activation of RIPK1 and RIPK3, as well as MLKL ^1, 22^. Our experiments confirm the hypothesis of parallel signaling cascades activated during necroptosis and reveal that LUBAC-dependent regulation affects both necroptotic NF-κB- and MLKL-dependent production of inflammatory cytokines. Apart from necroptotic signaling, necroptosis-independent functions of MLKL include the control of ICAM1, VCAM1 and E-selectin expression via RNA-binding motif protein 6 (RBM6) ^98^. Induction of necroptosis leads to a LUBAC- and MLKL-dependent upregulation of ICAM1 expression, suggesting that LUBAC also controls necroptosis-independent functions of MLKL. Taken together, these findings confirm parallel and independent LUBAC-dependent signaling events which most likely involve NF-κB and the core necroptotic machinery that converge downstream of activated MLKL to allow efficient membrane localization and necroptosis.

LUBAC inhibition prevents hotspot accumulation of phosphorylated MLKL as well as exocytosis in extracellular vesicles, suggesting that LUBAC-mediated M1 poly-Ub serves pro-necroptotic MLKL membrane thethering functions. Intruigingly, the M1 poly-Ub-dependent effects on MLKL mimic MLKL-specific monobodies that target the 4HB of human MLKL to inhibit necroptosis, but without affecting necrosome formation, MLKL activation and MLKL oligomerization ^99^. The functional role of the MLKL 4HB in M1 poly-Ub-mediated necroptosis inhibition remains to be investigated. Pharmacological interference of MLKL trafficking with inhibitors of Golgi, microtubule and actin networks (e.g. with Brefeldin A, Nocodazole and Cytochalasin A, respectively) also prevented membrane hotspot formation of activated MLKL ^21^. Intruigingly, loss of LUBAC triggered a similar phenotype, suggesting that LUBAC-mediated M1 poly-Ub might be involved in the regulation of MLKL trafficking.

Both endocytic and exocytic events balance MLKL membrane accumulation and involve the lipid raft-resided proteins FLOT1/2 ^53^, ESCRT complexes, ALIX, syntenin-1 and Rab27 ^22, 53, 74, 75, 100^. Recognition of ubiquitinated cargo by ESCRT complex subunits, like hepatocyte growth factor-regulated tyrosine kinase substrate (Hrs), signal-transducing adapter molecule (STAM) 1 and 2 and tumor susceptibility gene (TSG) 101 via ubiquitin-binding domains (UBDs), such as the Vps27/Hrs/Stam (VHS) domain, ubiquitin-interacting motif (UIM) domain and ubiquitin-conjugating enzyme E2 variant (UEV) domain, are important for endo- or exocytosis ^101–107^. Although the binding specificity and affinity of these UBD-containing proteins for M1 poly-Ub remains unclear, specific recognition of M1 poly-Ub might serve as docking platform to position activated MLKL in hotspots and to allow necroptosis. Intruigingly, FLOT1 is implicated in the transfer of ubiquitinated cargo between different ESCRT complexes by modulating the Ub-binding of Hrs upon epidermal growth factor (EGF) stimulation and release of Ub-mediated autoinhibition of the Hrs UIM domain ^108^. In line with this notion, we detected a strong necroptosis-induced and LUBAC-dependent enrichment of FLOT1/2 in M1 poly-Ub-based enrichments that was dependent on the activation of RIPK1, RIPK3 and MLKL. Additionally, loss of FLOT1/2 potentiated the HOIPIN-8-mediated decrease in necroptotic cell death. Although the necroptosis-induced increase in cellular M1 poly-Ub levels is not dependent on MLKL, inhibition of MLKL membrane translocation blocked the association of FLOT1 with M1 poly-Ub, indicating a direct link for FLOT1/2 in M1 poly-Ub and MLKL membrane localization and further demonstrating independent LUBAC-dependent signaling events during necroptosis.

Lipid rafts are important regulators of TNFα-mediated cell survival and death. Formation of the TNFR1 complex I in lipid rafts underlies RhoA and NF-κB activation, whereas activation of complex I outside lipid rafts promotes cell death ^109, 110^. Intruigingly, activated MLKL localizes to lipid rafts as well ^18^ and ALIX and ESCRT complexes regulate both MLKL endocytosis and TNFR1 internalization ^21, 111, 112^. As LUBAC and M1 poly-Ub are prominent components of the TNFR1-initiated complex I, a potential proximation of complex I, LUBAC, M1 poly-Ub and activated MLKL in lipid rafts upon TNFα-induced necroptosis, as well as shared signaling functions of M1 poly-Ub, cannot be excluded.

MLKL itself has been reported to be modified by mono-Ub upon MLKL oligomerization at membranes that might direct MLKL for proteasomal and lysosomal degradation ^113^. Intriguingly, although mutation of the identified mono-Ub sites in the MLKL 4HB did not prevent necroptosis, fusion of the non-specific DUB USP21 to MLKL did sensitize towards necroptosis ^113^. In addition, K63 poly-Ub of MLKL at K219 increased sensitivity towards necroptosis ^114^. Although the homologue residue in human MLKL, K230, is also ubiquitinated ^115^, it remains unclear if MLKL can be modified by M1 poly-Ub as well. Our findings further confirm that poly-Ub modification of MLKL or interacting proteins in close vicinity are important regulators of cell death.

We identified LUBAC-mediated M1 poly Ub as novel regulatory checkpoint that controls necroptosis in multicellular, primary human organoids *ex vivo*. The unique and first-time use of human primary organoids to systematically model necroptosis and the influence of LUBAC-mediated M1 poly-Ub in primary human organoids provides novel and relevant experimental models to study necroptosis in intact multicellular, near-physiological settings. Apart from necroptosis-proficient cell lines, experimental models to study necroptosis in human cells are extremely limited and so far no multicellular human models have been described. Since necroptosis underlies a wide variety of disease states, modeling necroptotic cell death in primary organoids will significantly contribute to gain new fundamental insights and to develop novel therapeutic options for emerging human diseases.

## Acknowledgments

The authors thank Laura Zein, Rebekka Karlowitz and all members of the van Wijk lab for advice, discussions and support during the study, Christina Hugenberg for proofreading and administrative support and Lorenza Lazzari (Unit of Regenerative Medicine, Cell Factory Fondazione IRCCS Ca’ Granda - Ospedale Maggiore Policlinico Milan, Italy) for providing the hPOs. Research in the lab of S.J.L.v.W. is supported by the Deutsche Forschungsgemeinschaft (DFG) (WI 5171/1-1, WI 5171/4-1, FU 436/20-1, FU 436/21-1 and project-ID 259130777 – CRC1177), the Deutsche Krebshilfe (70113680), Wilhelm Sander-Stiftung (2020.008.1), the Frankfurter Stiftung für krebskranke Kinder and the Dr. Eberhard and Hilde Rüdiger Foundation. Research in the group of F.P. is supported by EU FET-Open (828931), EU Transition Open (01 101057894), Wilhelm Sander-Stiftung (2020.008.1) and the German Aerospace Center (DLR) (IMMUNO3D-SHAPE). M.B. is supported by the DFG within the CRC1160 (Project ID 256073931-Z02), CRC/TRR167 (Project ID 259373024-Z01), CRC1453 (Project ID 431984000-S1), CRC1479 (Project ID: 441891347-S1). M.B. and G.A. also acknowledge funding from the German Federal Ministry of Education and Research (BMBF) within the Medical Informatics Funding Scheme - MIRACUM-FKZ 01ZZ1801B (M.B.) and EkoEstMed–FKZ 01ZZ2015 (G.A.).

## Author Contributions

N.W. and S.J.L.v.W. designed the study. N.W. performed most of the experiments. K.N.W. and F.P. performed experimentation on primary hPOs and analyzed the data. J.R. generated KO cell lines. S.S. and B.J. supported with experiments. L.F. and B.R. performed MACE-seq-based transcriptome analysis. G.A., T.M. and M.B. analyzed and visualized the MACE-seq data. N.W. and S.J.L.v.W. analyzed the data and prepared the manuscript. All authors have read, commented, and agreed on the submitted version of the manuscript.

## Materials and Methods

### Cell lines, reagents and chemicals

The human HT-29 adenocarcinoma cell line was obtained from DMSZ (Braunschweig, Germany) and the human acute monocytic leukemia cell line THP-1 was kindly provided by Thomas Oellerich (Frankfurt am Main, Germany) and authenticated by DSMZ. MEFs were kindly provided by Joanne Hildebrand (Melbourne, Australia). The L-929 mouse fibroblast cell line was obtained from ATCC (Manassas, VA, USA). HT-29 cells were cultured in McCoy’s 5A Medium +GlutaMAX^TM^-I (Thermo Fisher Scientific, Waltham, MA, USA) supplemented with 1% penicillin/streptomycin (Thermo Fisher Scientific) and 10% fetal bovine serum (FBS) (Thermo Fisher Scientific). THP-1 cells were maintained in RPMI1640 Medium +GlutaMAX^TM^-I (Thermo Fisher Scientific) supplemented with 1% sodium pyruvate (Thermo Fisher Scientific), 1% penicillin/streptomycin and 10% FBS. MEFs were cultured in DMEM +GlutaMAX^TM^-I (Thermo Fisher Scientific) supplemented with 1% sodium pyruvate, 1% penicillin/streptomycin and 10% FBS. L-929 cells were cultured in DMEM +GlutaMAX^TM^-I (Thermo Fisher Scientific) supplemented with 1% sodium pyruvate, 1% penicillin/streptomycin, 1% MEM Non-Essential Amino Acids (NEAA) Solution (Thermo Fisher Scientific) and 10% FBS. HT-29 MLKL KO, HT-29 RIPK3 KO and HT-29 RIPK3 KO cells re-expressing PAM-mutated Dox-inducible Strep-tagged WT RIPK3 have been described previously ^116^. All cell lines were cultured at 37°C with 5% carbon dioxide in a humidified atmosphere and regularly monitored for mycoplasma infection.

BV6 was kindly provided by Genentech Inc. (San Francisco, CA, USA), recombinant human TNFα was purchased from Biochrom Ltd. (Waterbeach, UK), Birinapant from Selleckchem (Houston, TX, USA), the pan-caspase inhibitors zVAD.fmk and Emricasan from Bachem AG (Bubendorf, Switzerland) and Selleckchem, Nec-1s, GSK’872 and NSA from Merck KGaA (Darmstadt, Germany), HOIPIN-8 from Axon Medchem LLC (Reston, VA, USA), Gliotoxin from Tocris Biosciences (Bristol, UK), Doxycycline hydrochloride from Merck KGgA (Darmstadt, Germany) and BafA1 from Selleckchem. All other chemicals were purchased from Carl Roth (Karlsruhe, Germany) or Merck KGaA, unless stated otherwise.

### Maintenance of primary hPOs

Primary hPOs are derived from the pancreatic duct of a male patient (age: 60, BMI: 26.3) and were provided by Lorenza Lazzari (Policlinico di Milano, Milan, Italy). The use of these hPOs is approved by the Institutional Review Board and the Pancreatic Islet Processing Unit, a National Transplant Center accredited facility (IT000679, https://webgate.ec.europa.eu/eucoding/reports/te/index.xhtml). hPOs were kept in an incubator at 37°C in a humidified atmosphere with 5% CO_2_, embedded in Cultrex Reduced Growth Factor BME, Type 2 (BME-2) droplets (R&D Systems, Minneapolis, MN, USA) and covered under medium. Medium was fully changed every 2 to 3 days and hPOs were passaged every 7 to 14 days depending on the proliferation rate and organoid size.

The organoid culture protocol was adapted from described procedures ^80^. Briefly, hPOs were maintained in basal medium, composed of Advanced DMEM/F-12 (Thermo Fisher Scientific) with 1x GlutaMAX™ Supplement (Thermo Fisher Scientific), 100 U/mL penicillin/streptomycin (Thermo Fisher Scientific) and 10 mM HEPES (Thermo Fisher Scientific), supplemented with 1x B-27™ (minus vitamin A, Thermo Fisher Scientific), 1x N-2 (Thermo Fisher Scientific), 2,5 mM *N*-acetylcysteine (MilliporeSigma, St. Louis, US), 1 µg/mL recombinant human R-Spondin1 (PeproTech, Rocky Hill, US), 10 mM nicotinamide (MilliporeSigma), 50 ng/mL recombinant human EGF (PeproTech), 100 ng/mL recombinant human FGF-10 (PeproTech), 25 ng/mL Noggin (PeproTech), 10 nM [Leu^15^]-Gastrin I (MilliporeSigma), 5 µM A83-01 (Tocris Bioscience, Minneapolis, US), 10 µM Forskolin (Tocris Bioscience) and 3 µM Prostaglandin E_2_ (Tocris Bioscience) (hPO medium).

For passaging of the primary hPOs, BME droplets were disrupted by pipetting and organoids were retrieved by centrifugation in ice-cold hPO medium. Organoids were sheared into fragments prior to resuspension in ice-cold liquified BME-2 and seeded as droplets in pre-warmed non-treated CELLSTAR® 48-well suspension culture plates (Greiner Bio-One GmbH, Kremsmünster, AUT). Organoid-containing BME-2 droplets were polymerized for 15 min at 37°C after which the droplets were overlaid with hPO medium.

### Induction and quantification of necroptotic cell death

The indicated cells were seeded in CELLSTAR® 96-well tissue culture plates (Greiner Bio-One GmbH) at appropriate densities the day before treatment. For necroptosis induction, cells were pretreated with 1 µM BV6 and 20 µM zVAD.fmk for 30 min (if not stated otherwise), followed by stimulation with 10 ng/mL recombinant human TNFα for the indicated time (TBZ). The inhibitors HOIPIN-8 (30 µM), Gliotoxin (1 µM), Nec-1s (30 µM), GSK’872 (20 µM) and NSA (10 µM) were added 1 h prior to TBZ treatment. Cell death was determined by PI and Hoechst33342 (Hoe) staining and fluorescence-based quantification of dead (PI-positive) cells compared to the total cell number (Hoe-positive cells) using an ImageXpress® Micro XLS Widefield High-Content Analysis System and MetaXpress® Software (Molecular Devices Sunnyvale, CA, USA).

### Induction and imaging of necroptotic cell death in primary hPOs

hPOs were seeded in 5 µL BME-2 droplets in sterile CELLSTAR® 96-well suspension culture plates (Greiner Bio-One GmbH) and overlaid with 100 µL hPO medium. hPOs were cultured for 3-5 days prior to experiments to assure organoid formation and growth. For pre-treatment with HOIPIN-8, hPO medium was replaced by hPO medium with the indicated concentrations of HOIPIN-8 and hPOs were placed back in the incubator. After 30 min, hPO medium was again replaced with medium supplemented with the indicated treatments and the plates were transferred to the Zeiss AxioObserver Z1 microscope. Incubation was done at 37°C and 5% CO_2_ during which 24 h time lapse imaging in 30 min intervals was performed. Each well was imaged as tiles of two x two with eleven Z-slices of 60 µm thickness.

For live/dead assays, primary hPOs were stained using 10 µg/mL PI (Millipore Sigma), 0.5 µg/mL FDA (Millipore Sigma) and 200 µg/mL Hoe (ThermoFisher Scientific) for 30 min. Dyes were removed and the wells were washed with pre-warmed PBS prior to imaging in PBS, using the same microscope settings as described above.

Images were exported using the ZEN 2.6 lite software (Carl Zeiss Microscopy GmbH, Oberkochen, Germany) and subsequently processed using the FiJi/ImageJ software.

### RNAi-mediated silencing

Transient genetic silencing of HOIP and FLOT1/2 was performed by reverse transfection of HT-29 cells with the following siRNAs: human siHOIP (RNF31) (Silencer® Select siRNA #1 s30108, #2 s30109, #3 s30110, 40 nM) (Thermo Fisher Scientific, Waltham, MA, USA), human FLOT1 (ON-TARGETplus Human FLOT1 siRNA SMART POOL, L-010636-00-0010) (Horizon Discovery Ltd., Waterbeach, UK), human FLOT2 (ON-TARGETplus Human FLOT2 siRNA SMART POOL, L-003666-01-0010) (Horizon Discovery Ltd.), non-targeting control siRNA (#4390843, 40 nM) (Thermo Fisher Scientific). In brief, the indicated cell lines were prepared in penicillin/streptomycin-free medium and combined with the siRNA-lipid complexes, prepared in OptiMEM (Thermo Fisher Scientific) using Lipofectamine™ RNAiMAX Transfection Reagent (Thermo Fisher Scientific) according to the manufacturer’s protocol and incubated for 5 min at room temperature. The medium was changed to full medium after 6 h and the cells were reseeded for the experiments 48 h post-transfection.

### Western blotting

For general cell lysis, RIPA lysis buffer (50 mM Tris-HCl pH 8, 150 mM NaCl, 1% Nonidet P-40 (NP-40), 2 mM MgCl_2_, 0.5% sodium deoxycholate) supplemented with protease inhibitor complex (PIC) (Roche, Grenzach, Germany), 0.1% SDS, 1 mM sodium orthovanadate, 5 mM sodium fluoride, 1 mM β-gylcerophosphate, 1 mM phenylmethylsulfonyl fluoride and Pierce Universal Nuclease (Thermo Fisher Scientific) was used. Indicated cell lines were incubated with lysis buffer for 30 min on ice. The lysates were cleared by centrifugation at 18,000 x *g* and 4°C for 20 min and protein concentration was determined using the BCA Protein Assay Kit from Pierce™ (Thermo Fisher Scientific) according to the manufacturer’s protocol. Approximately 30-50 µg protein were boiled in 6x SDS loading buffer (350 mM Tris Base pH 6.8, 38% glycerol, 10% SDS, 93 mg/ml dithiothreitol (DTT), 120 mg/ml bromophenol blue) followed by Western blot analysis. For Western blot analysis of MLKL oligomers, indicated cell lines were lysed as described above in 1% NP-40 lysis buffer (20 mM Tris-HCl pH 7.5, 50 mM NaCl, 5 mM EDTA, 10% glycerol, 1% NP-40) supplemented with PIC, 1 mM sodium orthovanadate, 5 mM sodium fluoride, 1 mM β-gylcerophosphate and Pierce Universal Nuclease. The cleared lysates were boiled in 6x SDS loading buffer or non-reducing 6x SDS loading buffer (350 mM Tris Base pH 6.8, 38% glycerol, 10% SDS, 120 mg/ml bromophenol blue), followed by Western blot analysis. The following antibodies were used in this study: human anti-M1 Ub (Genentech, San Francisco, CA, USA), rabbit anti-RNF31/HOIP (ab46322, Abcam, Cambridge, UK), mouse anti-HOIL-1 (MABC576, Merck KGaA, Darmstadt, Germany), rabbit anti-Sharpin (12541S, Cell Signaling Technology, Danvers, MA, USA), mouse anti-phospho-IκBα (S32/S36) (9246L, Cell Signaling Technology), rabbit anti-IκBα (9242S, Cell Signaling Technology), rabbit anti-phospho-RIPK1 (S166) (65746S, Cell Signaling Technology), rabbit anti-phospho-RIPK1 (S166) (31122S, Cell Signaling Technology), mouse anti-RIPK1 (551041, BD Biosciences, San Jose, CA, USA), mouse anti-RIPK1 (610459, BD Biosciences), rabbit anti-phospho-RIPK3 (S277) (ab209384, Abcam), rabbit anti-RIPK3 (13526S, Cell Signaling Technology), rabbit anti-phospho-MLKL (S358) (91689S, Cell Signaling Technology), rabbit anti-phospho-MLKL (S345) (ab196436, Abcam), rabbit anti-MLKL (14993S, Cell Signaling Technology), rabbit anti-MLKL (orb32399, Biorbyt Ltd., Cambridge, UK), mouse anti-ALIX (634502, BioLegend, San Diego, CA, USA), mouse anti-CD9 (sc-13118, Santa Cruz Biotechnologies, Santa Cruz, CA, USA), mouse anti-Flotillin-1 (610820, BD Biosciences), mouse anti-Flotillin-2 (610383, BD Biosciences), mouse anti-GAPDH (5G4cc, HyTest Ltd, Turku, Finland), mouse anti-Vinculin (V9131, Merck KGaA), mouse anti-β-actin (A5441, Merck KGaA). The following horseradish peroxidase (HRP)-coupled secondary antibodies were used for detection using Pierce™ ECL Western Blotting-Substrate (Thermo Fisher Scientific): HRP-conjugated goat anti-mouse IgG (ab6789, Abcam), HRP-conjugated goat anti-rabbit IgG (ab6721, Abcam), HRP-conjugated goat anti-human IgG (AP309P, Merck KGaA). For stripping and re-probing of Western blot membranes, membranes were incubated with 0.4 M NaOH solution followed by blocking for 1 h. Representative blots of at least two independent experiments are shown. If the samples of one experiment are detected on multiple Western blotting membranes only one representative loading control is shown for clarity.

### RIPK3 immunoprecipitation

For RIPK3 immunoprecipitations, HT-29 RIPK3 KO cells re-expressing PAM-mutated Dox-inducible Strep-tagged RIPK3 WT ^116^ were seeded in full medium and RIPK3 expression was induced by adding 1 µg/mL Dox 6 h after seeding. Prior to treatment, the medium was changed to fresh medium and the cells were lysed in 1% NP-40 lysis buffer after treatment (as described above). For Strep-RIPK3 immunoprecipitations, MagStrep “type3” XT beads (IBA Lifesciences GmbH, Göttingen, Germany) were pre-washed twice in 1% NP-40 lysis buffer and incubated with 1,500-1,800 µg protein over night at 4°C on a rotating wheel. On the next day, the beads were washed five times with TBS-T (20 mM Tris, 150 mM NaCl, 0.1%Tween 20, pH 8.0) using a magnetic separator and then boiled in 2x SDS loading buffer for 5 min at 96°C followed by Western blotting.

### UBAN pulldowns

M1 ubiquitinated proteins were enriched using GST-tagged UBAN-coupled Glutathione MagBeads (Genscript, Piscataway NJ, SA), as described previously ^116^. Briefly, cell lysates were prepared in 1% NP-40 lysis buffer (as described above). 1,800-2,000 µg protein were incubated with pre-washed (NP-40 lysis buffer) GST-UBAN beads overnight on a rotating wheel at 4°C. On the following day, the beads were washed five times with TBS-T using a magnetic separator, then boiled in 2x SDS loading buffer for 5 min at 96°C and analyzed by Western blotting. Equal loading of the GST-UBAN beads was confirmed using ponceau S (Merck KGaA) staining.

### Isolation of exosomes

For the enrichment of exosomes, the indicated HT-29 cell lines were seeded in full medium the day before treatment. Prior to the addition of compounds, the medium was removed, cells were washed with PBS and fresh FBS-free medium was added. After treatment, the cell culture supernatant was collected for exosome isolation and cells were lysed in RIPA lysis buffer as described above. The supernatant was cleared by centrifugation (2,000 x *g*, 30 min 4°C) in order to remove cell debris. Exosome isolation was performed using the Total Exosome Isolation Reagent (from cell culture media) (Thermo Fisher Scientific) according to the manufacturer’s instructions. The cleared lysates were mixed with 0.5 volumes of the Total Exosome Isolation reagent and incubated over night at 4°C. The next day, the samples were centrifuged for 1 h at 10,000 x *g* at 4°C. The supernatants were discarded and the pelleted exosomes were lysed in RIPA lysis buffer, followed by incubation in an appropriate amount of 6x SDS loading buffer at 96°C for 5 min. Exosome lysates and cell lysates were analyzed by Western blotting.

### Cell fractionation

Cell fractionation by phase separation using Triton X-114 lysis buffer was performed as described before ^17^. In brief, indicated cell lines were treated and lysed in Triton-X114 (MilliporeSigma) lysis buffer (150 mM NaCl, 20 mM HEPES pH 7.4, 2% Triton X-114), supplemented with PIC, 1 mM sodium orthovanadate, 5 mM sodium fluoride and 1 mM β-gylcerophosphate, for 30 min on ice, followed by 10 min centrifugation at 15,000 x *g* at 4°C. The supernatant was warmed at 30°C for 4 min and centrifuged at 1,500 x *g* for 5 min. The aqueous layer of the two-phase solution was collected and re-centrifuged at 1,500 x *g* for 5 min to remove contaminations from the detergent-enriched phase. The detergent-enriched fraction was washed twice with basal buffer (150 mM NaCl, 20 mM HEPES pH 7.4) by centrifugation 1,500 x *g* for 5 min and then diluted in basal buffer. The protein concentrations of both the aqueous and detergent-enriched fractions were determined and 40-50 µg of protein were incubated in an appropriate amount of 6x SDS loading buffer at 96°C for 5 min, followed by Western blotting.

### RNA isolation, cDNA synthesis and quantitative real-time PCR

Total RNA was prepared using the peqGOLD total RNA isolation kit (VWR, Radnor, PA, USA) according to manufacturer’s protocol. In brief, cell lysates were prepared using RNA lysis buffer, transferred to peqGOLD RNA Homogenizer Columns and centrifuged at 13,000 x *g* for 1 min. The filtrates were mixed with an equal volume of 70% ethanol and loaded to a peqGOLD RNA Mini Column followed by centrifugation at 10,000 x*g* for 1 min. After a washing step with RNA Wash Buffer I and two washing steps with 80% ethanol (centrifugation at 10,000 x *g*), the columns were dried by centrifugation at 12,000 x *g* for 2 min. The RNA was eluted in nuclease-free water by centrifugation at 12,000 x *g* for 2 min. For cDNA synthesis, 1 µg of RNA was transcribed using the RevertAid H Minus First Strand Kit (ThermoFisher Scientific) according to the manufacturer’s protocol. CXCL1, CXCL10, TNFα and ICAM1 gene expression levels were determined using SYBR green-based quantitative real-time PCR (Applied Biosystems, Darmstadt, Germany) using the 7900GR Fast Real-time PCR system (Applied Biosystems). Data were normalized against 18S-rRNA and RPII expression and relative gene expression levels were calculated using the ΔΔCt-method. The primers used in this study are listed below and were obtained from Eurofins (Hamburg, Germany). CXCL1: forward AACCGAAGTCATAGCCACAC, reverse GTTGGATTTGTCACTGTTCAGC. CXCL10: forward CTGAGCCTACAGCAGAGGAAC, reverse GATGCAGGTACAGCGTACAGT. TNFα: forward ACAACCCTCAGACGCCACAT, reverse TCCTTTCCAGGGGAGAGAGG. ICAM1: forward CTTCCTCACCGTGTACTGGAC, reverse GGCAGCGTAGGGTAAGGTTC. 28S-rRNA: forward TTGAAAATCCGGGGGAGAG, reverse ACATTGTTCCAACATGCCAG, RPII: forward GCACCACGTCCAATGACAT, reverse GTGCGGCTGCTTCCATAA.

### Quantification of CXCL1 cytokine secretion

To determine CXCL1 secretion from necroptotic cells, the indicated cell lines were seeded in 96-well dishes and treated as described above. After treatment, supernatants were collected and cleared by centrifugation (300 x *g*, 5 min, 4°C). CXCL1 levels were measured after 3 h and 6 h using the Human CXCL1/GROα DuoSet® ELISA kit (R&D Systems, Minneapolis, MN, USA) according to the manufacturer’s instructions. Absorbance at 450 nm was determined using the Infinite M200 microplate reader (Tecan, Crailsheim, Germany).

### Immunofluorescence staining

For immunofluorescence staining of phosphorylated MLKL, indicated cell lines were seeded in CELLSTAR® 96-well tissue culture plates (Greiner Bio-One GmbH) at an appropriate density the day before treatment. After treatment, the supernatant was removed and the cells were fixed by incubation in 4% paraformaldehyde solution for 20 min at room temperature, followed by permeabilization for 15 min with 0.1% Triton X-100. After three washing steps with PBS, the cells were blocked in antibody dilution buffer (ADB) (0.9% NaCl, 10 mM Tris/HCl pH 7.5, 5 mM EDTA, 1 mg/mL BSA) for 45 min, followed by incubation with rabbit anti-phospho-MLKL (S358) (1:100, ab187091, Abcam) (diluted in ADB) overnight at 4°C. On the next day, the cells were washed five times with PBS and incubated with Cy3 AffiniPure Donkey Anti-Rabbit IgG (H+L) (1:800, 711-165-152, Jackson Immuno Research, West Grove, PA, USA) and 0.1 µg/mL DAPI (diluted in ADB) for 90 min at room temperature. After five washing steps with PBS the cells were analyzed using the ImageXpress® Micro XLS Widefield High-Content Analysis System and MetaXpress® Software (Molecular Devices).

### Massive Analysis of cDNA Ends (MACE-seq)

Massive Analyses of cDNA Ends (MACE-Seq) was performed by GenXPro GmbH in Frankfurt am Main using the MACE-Seq kit according to the manual of the manufacturers. Briefly, cDNA was generated from fragmented RNA with barcoded poly-A primers during reverse transcription. After second-strand synthesis and 5’ adapter integration, a PCR with minimum number of cylces was used to produce a library that was sequenced on an Illumina NextSeq500 machine with 1x 75 bps ^117^.

### MACE-seq analysis

Raw MACE-seq data was preprocessed using Cutadapt ^118^ to eliminate poly-A-tails as well as bad-quality base reads. FastQC was used to assess the quality of sequencing after trimming. Cleaned reads were mapped and quantified with a reference genome using STAR. ENSEMBL-GTF data were used to provide genomic locations for quantification as well as additional data for annotation (such as gene name, gene description, Gene Ontology (GO)-Terms etc). Differential expression analysis was performed using DESeq2 ^119^, which is based on negative binomial generalized linear models. Results were compiled into a final table including significance parameters (pvalue, FDR) and log2FoldChanges. BigWig track files were generated by deepTools. Final data visualization of the significantly expressed and the differentially regulated genes was performed by custom R-Scripts.

### Statistical analysis

Statistical significance was determined with the GraphPad Prism 9 Software (GraphPad Software, La Jolla CA, USA) using 2-way ANOVA followed by Tukey’s multiple comparisons test. *P* values < 0.05 are considered significant and depicted as follows: *P* ≤ 0.05: *, *P* ≤ 0.01: **, *P* ≤ 0.001: ***, *P* ≤ 0.0001: ****, ns: not significant.

## Data availability

MACE-seq data have been deposited at GEO and are publicly available as of the date of publication. All primary data will be shared by the lead contact upon request.

## Supplemental Information

**Figure S1. Related to Figure 1. A.** Quantification of cell death in untreated (UT) and Gliotoxin (1 µM)-pre-treated HT-29 cells after treatment with TBZ (10 ng/mL TNFα, 1 µM BV6, 20 µM zVAD.fmk) for the indicated time points. Mean and SEM of at least three independent experiments are shown. \*\*\*\**P*<0.0001. **B.** Quantification of cell death in untreated (UT) and Gliotoxin (1 µM)-pre-treated THP-1 cells after treatment with TBZ (10 ng/mL TNFα, 1 µM BV6, 20 µM zVAD.fmk) for the indicated time points. Mean and SEM of at least three independent experiments are shown. \*\*\**P*<0.001; \*\*\*\**P*<0.0001. **C.** Western blot analysis of HOIP levels in control (siCtrl) and HOIP knockdown (siHOIP) HT-29 cells. GAPDH was used as loading control. Representative blots of at least two independent experiments are shown. Related to Figure 1C. **D.** Western blot analysis of total M1 Ub levels in control or HOIPIN-8 (30 µM)-pre-treated THP-1 cells treated with TBZ (10 ng/mL TNFα, 1 µM BV6, 20 µM zVAD.fmk) for 4 h. GAPDH was used as loading control. Representative blots of at least two independent experiments are shown. **E.** Amino acid sequence alignment showing conservation of human (aa841-aa960) and murine HOIP (RNF31) (aa835-aa954). Conserved residues interacting with HOIPIN-8 are highlighted and indicated by arrowheads. Sequence alignment was performed using the CLUSTAL O (1.2.4) algorithm. **F.** Western blot analysis of phosphorylated and total IκBα levels in control or HOIPIN-8 (30 µM)-pre-treated HT-29 cells upon treatment with 10 ng/mL TNFα for the indicated time points. Vinculin was used as loading control. Representative blots of at least two independent experiments are shown. **G.** Western blot analysis of phosphorylated and total IκBα levels in control or HOIPIN-8 (30 µM)-pre-treated MEFs upon treatment with 10 ng/mL TNFα for the indicated time. Vinculin was used as loading control. Representative blots of at least two independent experiments are shown. **H.** Quantification of cell death in untreated (UT) and HOIPIN-8 (30 µM) pre-treated MEFs after treatment with TBZ (10 ng/mL TNFα, 1 µM BV6, 20 µM zVAD.fmk) for the indicated time points. Mean and SEM of at least three independent experiments are shown. \*\*\**P*<0.001; \*\*\*\**P*<0.0001; ns not significant. **I.** Quantification of cell death in untreated (UT) and HOIPIN-8 (30 µM)-pre-treated, GSK’872 (20 µM)-pre-treated or HOIPIN-8 and GSK’872-pre-treated L-929 cells after treatment with TBZ (10 ng/mL TNFα, 1 µM BV6, 20 µM zVAD.fmk) for the indicated time points. Mean and SEM of at least three independent experiments are shown. \*\*\*\**P*<0.0001; ns not significant. **J.** Western blot analysis of total M1 Ub levels in control or HOIPIN-8 (30 µM)-pre-treated MEFs treated with TBZ (10 ng/mL TNFα, 1 µM BV6, 20 µM zVAD.fmk) for 3 h. GAPDH was used as loading control. Representative blots of at least two independent experiments are shown.

**Figure S2. Related to Figure 2. A.** Western blot analysis of LUBAC subunits HOIP and HOIL-1 expression levels in control *versus* HOIPIN-8 (30 µM)-pre-treated MEFs upon treatment with TBZ (10 ng/mL TNFα, 1 µM BV6, 20 µM zVAD.fmk) for 3 h. GAPDH was used as loading control. Representative blots of at least two independent experiments are shown. **B.** Western blot analysis of expression levels of phosphorylated and total RIPK1 and MLKL upon 4 h TBZ (10 ng/mL TNFα, 1 µM BV6, 20 µM zVAD.fmk) treatment of THP-1 cells pre-treated with HOIPIN-8 (30 µM). Vinculin was used as loading control. Representative blots of at least two independent experiments are shown. **C.** Western blot analysis of expression levels of phosphorylated and total RIPK1, RIPK3 and MLKL, and of M1 Ub in TBZ-treated (10 ng/mL TNFα, 1 µM BV6, 20 µM zVAD.fmk) control (nht), RIPK3 KO and MLKL KO HT-29 cells pre-treated with HOIPIN-8 (30 µM). GAPDH was used as loading control. Representative blots of at least two independent experiments are shown.

**Figure S3. Related to Figure 3. A.** mRNA expression levels of CXCL1 and CXCL10 of control (nht) or MLKL KO HT-29 cells treated with TBZ (10 ng/mL TNFα, 1 µM BV6, 20 µM zVAD.fmk) for 3 h. Gene expression was normalized against 18S and RPII mRNA expression and is presented as x-fold mRNA expression compared to the untreated (UT) control cells. Mean and SEM of at least three independent experiments are shown. \*\**P*<0.01; \*\*\**P*<0.001. **B.** mRNA expression levels of TNFα of control (UT), HOIPIN-8 (30 µM) or NSA (10 µM)-pre-treated HT-29 cells upon treatment with TBZ (10 ng/mL TNFα, 1 µM BV6, 20 µM zVAD.fmk) for 3 h. Gene expression was normalized against 18S and RPII mRNA expression and is presented as x-fold mRNA expression compared to the untreated (UT) control. Mean and SEM of at least three independent experiments are shown. \*\*\*\**P*<0.0001. **C.** mRNA expression levels of ICAM1 of control (UT), HOIPIN-8 (30 µM) or NSA (10 µM)-pre-treated HT-29 cells upon treatment with TBZ (10 ng/mL TNFα, 1 µM BV6, 20 µM zVAD.fmk) for 3 h. Gene expression was normalized against 18S and RPII mRNA expression and is presented as x-fold mRNA expression compared to the untreated (UT) control. Mean and SEM of at least three independent experiments are shown. \*\*\*\**P*<0.0001.

**Figure S4. Related to Figure 4. A.** Distribution of M1 poly-Ub in the micelle-poor (aqueous) and micelle-rich (detergent) fraction after phase separation using Triton X-114 lysis buffer in untreated (UT) and HOIPIN-8 (30 µM)-pre-treated HT-29 cells upon treatment with TBZ (10 ng/mL TNFα, 1 µM BV6, 20 µM zVAD.fmk) for 4 h. GAPDH was used as loading control for soluble proteins and CD9 as loading control for membrane proteins. Representative blots of at least two independent experiments are shown. **B.** Fluorescence microscopy images show localization of phosphorylated MLKL (green) in untreated (UT) and NSA (10 µM)-pre-treated HT-29 cells and after treatment with TBZ (10 ng/mL TNFα, 1 µM BV6, 20 µM zVAD.fmk) for 3 h. Nuclei were stained with DAPI (blue). Representative images of at least two independent experiments are shown. Scale bar 50 µm.

**Figure S5. Related to Figure 5. A.** GST-UBAN-mediated pulldown of M1 ubiquitinated proteins in control or HOIPIN-8 (30 µM)-pre-treated THP-1 cells treated with TBZ (10 ng/mL TNFα, 1 µM BV6, 20 µM zVAD.fmk) for 4 h. GST and GAPDH were used as loading controls. Representative blots of at least two independent experiments are shown. **B.** Western blot analysis of FLOT1/2 levels in control (siCtrl) and FLOT1 (siFLOT1), FLOT2 (siFLOT2) or combined FLOT1/2 (siFLOT1+2) knockdown HT-29 cells. Vinculin was used as loading control. Representative blots of at least two independent experiments are shown. Related to Figure 5B. **C.** Western blot analysis of total M1 Ub levels of control or HOIPIN-8 (30 µM)-pre-treated control (siCtrl) or FLOT1/2 (siFLOT1+2) knockdown HT-29 cells after treatment with TBZ (10 ng/mL TNFα, 1 µM BV6, 20 µM zVAD.fmk) for 4 h. GAPDH was used as loading control. Representative blots of at least two independent experiments are shown.

**Figure S6. Related to Figure 6. A.** Representative images of time-lapse videos of untreated (UT), Birinapant (20 µM) and Emricasan (10 µM) (BiE)-treated, HOIPIN-8 (30 µM)-pre-treated and BiE-treated (BiE+HOIPIN-8) and NSA (10 µM)-pre-treated and BiE-treated (BiE+NSA) primary hPOs for a total period of 23.5 h. Scalebars: 250 µm. **B.** *Idem* as A., but treated primary hPOs were stained with Hoechst33342 (Hoe) (blue), FDA (live; green) and PI (dead; red) after 24 h treatment with BiE, BiE+HOIPIN-8 or BiE+NSA prior to imaging. Scalebars: 250 µm.

**Figure S7. Related to Figure 6.** Representative images of untreated (UT), BV6 (5 µM) and zVAD.fmk (20 µM) (BZ)-treated and HOIPIN-8 (30 µM)-pre-treated and BZ-treated (BZ+HOIPIN-8) primary hPOs that were stained with Hoechst33342 (Hoe) (blue), FDA (live; green) and PI (dead; red) after 24 h treatment prior to imaging. Scalebars: 250 µm.

**Movie 1. Related to Figure 6A.** Time-lapse videos of untreated (UT), BV6 and Emricasan (BE)-treated and HOIPIN-8-pre-treated and BE (BE+HOIPIN-8)-treated primary hPOs for a total period of 23.5 h. Scalebars: 250 µm.

**Movie 2. Related to Figure S6A.** Time-lapse videos of untreated (UT), Birinapant and Emricasan (BiE)-treated, HOIPIN-8-pre-treated and BiE (BiE+HOIPIN-8)-treated and NSA-pre-treated and BiE (BiE+NSA)-treated primary hPOs for a total period of 23.5 h. Scalebars: 250 µm.

## References

1. Zhu, K., Liang, W., Ma, Z., Xu, D., Cao, S., Lu, X., Liu, N., Shan, B., Qian, L., and Yuan, J. (2018). Necroptosis promotes cell-autonomous activation of proinflammatory cytokine gene expression. Cell Death Dis 9, 500. 10.1038/s41419-018-0524-y.

2. Kaczmarek, A., Vandenabeele, P., and Krysko, D.V. (2013). Necroptosis: the release of damage-associated molecular patterns and its physiological relevance. Immunity 38, 209–223. 10.1016/j.immuni.2013.02.003.

3. Weinlich, R., Oberst, A., Beere, H.M., and Green, D.R. (2017). Necroptosis in development, inflammation and disease. Nat Rev Mol Cell Biol 18, 127–136. 10.1038/nrm.2016.149.

4. Pasparakis, M., and Vandenabeele, P. (2015). Necroptosis and its role in inflammation. Nature 517, 311–320. 10.1038/nature14191.

5. Emmerich, C.H., Bakshi, S., Kelsall, I.R., Ortiz-Guerrero, J., Shpiro, N., and Cohen, P. (2016). Lys63/Met1-hybrid ubiquitin chains are commonly formed during the activation of innate immune signalling. Biochem Biophys Res Commun 474, 452–461. 10.1016/j.bbrc.2016.04.141.

6. Emmerich, C.H., Ordureau, A., Strickson, S., Arthur, J.S., Pedrioli, P.G., Komander, D., and Cohen, P. (2013). Activation of the canonical IKK complex by K63/M1-linked hybrid ubiquitin chains. Proc Natl Acad Sci U S A 110, 15247–15252. 10.1073/pnas.1314715110.

7. Bertrand, M.J., Milutinovic, S., Dickson, K.M., Ho, W.C., Boudreault, A., Durkin, J., Gillard, J.W., Jaquith, J.B., Morris, S.J., and Barker, P.A. (2008). cIAP1 and cIAP2 facilitate cancer cell survival by functioning as E3 ligases that promote RIP1 ubiquitination. Mol Cell 30, 689–700. 10.1016/j.molcel.2008.05.014.

8. Draber, P., Kupka, S., Reichert, M., Draberova, H., Lafont, E., de Miguel, D., Spilgies, L., Surinova, S., Taraborrelli, L., Hartwig, T., et al. (2015). LUBAC-Recruited CYLD and A20 Regulate Gene Activation and Cell Death by Exerting Opposing Effects on Linear Ubiquitin in Signaling Complexes. Cell Rep 13, 2258–2272. 10.1016/j.celrep.2015.11.009.

9. Wertz, I.E., Newton, K., Seshasayee, D., Kusam, S., Lam, C., Zhang, J., Popovych, N., Helgason, E., Schoeffler, A., Jeet, S., et al. (2015). Phosphorylation and linear ubiquitin direct A20 inhibition of inflammation. Nature 528, 370–375. 10.1038/nature16165.

10. Mompean, M., Li, W., Li, J., Laage, S., Siemer, A.B., Bozkurt, G., Wu, H., and McDermott, A.E. (2018). The Structure of the Necrosome RIPK1-RIPK3 Core, a Human Hetero-Amyloid Signaling Complex. Cell 173, 1244–1253 e1210. 10.1016/j.cell.2018.03.032.

11. Li, J., McQuade, T., Siemer, A.B., Napetschnig, J., Moriwaki, K., Hsiao, Y.S., Damko, E., Moquin, D., Walz, T., McDermott, A., et al. (2012). The RIP1/RIP3 necrosome forms a functional amyloid signaling complex required for programmed necrosis. Cell 150, 339–350. 10.1016/j.cell.2012.06.019.

12. Cho, Y.S., Challa, S., Moquin, D., Genga, R., Ray, T.D., Guildford, M., and Chan, F.K. (2009). Phosphorylation-driven assembly of the RIP1-RIP3 complex regulates programmed necrosis and virus-induced inflammation. Cell 137, 1112–1123. 10.1016/j.cell.2009.05.037.

13. He, S., Wang, L., Miao, L., Wang, T., Du, F., Zhao, L., and Wang, X. (2009). Receptor interacting protein kinase-3 determines cellular necrotic response to TNF-alpha. Cell 137, 1100–1111. 10.1016/j.cell.2009.05.021.

14. Sun, L., Wang, H., Wang, Z., He, S., Chen, S., Liao, D., Wang, L., Yan, J., Liu, W., Lei, X., and Wang, X. (2012). Mixed lineage kinase domain-like protein mediates necrosis signaling downstream of RIP3 kinase. Cell 148, 213–227. 10.1016/j.cell.2011.11.031.

15. Murphy, J.M., Czabotar, P.E., Hildebrand, J.M., Lucet, I.S., Zhang, J.G., Alvarez-Diaz, S., Lewis, R., Lalaoui, N., Metcalf, D., Webb, A.I., et al. (2013). The pseudokinase MLKL mediates necroptosis via a molecular switch mechanism. Immunity 39, 443–453. 10.1016/j.immuni.2013.06.018.

16. Su, L., Quade, B., Wang, H., Sun, L., Wang, X., and Rizo, J. (2014). A plug release mechanism for membrane permeation by MLKL. Structure 22, 1489–1500. 10.1016/j.str.2014.07.014.

17. Wang, H., Sun, L., Su, L., Rizo, J., Liu, L., Wang, L.F., Wang, F.S., and Wang, X. (2014). Mixed lineage kinase domain-like protein MLKL causes necrotic membrane disruption upon phosphorylation by RIP3. Mol Cell 54, 133–146. 10.1016/j.molcel.2014.03.003.

18. Chen, X., Li, W., Ren, J., Huang, D., He, W.T., Song, Y., Yang, C., Li, W., Zheng, X., Chen, P., and Han, J. (2014). Translocation of mixed lineage kinase domain-like protein to plasma membrane leads to necrotic cell death. Cell Res 24, 105–121. 10.1038/cr.2013.171.

19. Hildebrand, J.M., Tanzer, M.C., Lucet, I.S., Young, S.N., Spall, S.K., Sharma, P., Pierotti, C., Garnier, J.M., Dobson, R.C., Webb, A.I., et al. (2014). Activation of the pseudokinase MLKL unleashes the four-helix bundle domain to induce membrane localization and necroptotic cell death. Proc Natl Acad Sci U S A 111, 15072–15077. 10.1073/pnas.1408987111.

20. Dondelinger, Y., Declercq, W., Montessuit, S., Roelandt, R., Goncalves, A., Bruggeman, I., Hulpiau, P., Weber, K., Sehon, C.A., Marquis, R.W., et al. (2014). MLKL compromises plasma membrane integrity by binding to phosphatidylinositol phosphates. Cell Rep 7, 971–981. 10.1016/j.celrep.2014.04.026.

21. Samson, A.L., Zhang, Y., Geoghegan, N.D., Gavin, X.J., Davies, K.A., Mlodzianoski, M.J., Whitehead, L.W., Frank, D., Garnish, S.E., Fitzgibbon, C., et al. (2020). MLKL trafficking and accumulation at the plasma membrane control the kinetics and threshold for necroptosis. Nat Commun 11, 3151. 10.1038/s41467-020-16887-1.

22. Gong, Y.N., Guy, C., Olauson, H., Becker, J.U., Yang, M., Fitzgerald, P., Linkermann, A., and Green, D.R. (2017). ESCRT-III Acts Downstream of MLKL to Regulate Necroptotic Cell Death and Its Consequences. Cell 169, 286–300 e216. 10.1016/j.cell.2017.03.020.

23. Cai, Z., Jitkaew, S., Zhao, J., Chiang, H.C., Choksi, S., Liu, J., Ward, Y., Wu, L.G., and Liu, Z.G. (2014). Plasma membrane translocation of trimerized MLKL protein is required for TNF-induced necroptosis. Nat Cell Biol 16, 55–65. 10.1038/ncb2883.

24. Seo, J., Nam, Y.W., Kim, S., Oh, D.B., and Song, J. (2021). Necroptosis molecular mechanisms: Recent findings regarding novel necroptosis regulators. Exp Mol Med 53, 1007–1017. 10.1038/s12276-021-00634-7.

25. Karlowitz, R., and van Wijk, S.J.L. (2021). Surviving death: emerging concepts of RIPK3 and MLKL ubiquitination in the regulation of necroptosis. FEBS J. 10.1111/febs.16255.

26. Samson, A.L., Garnish, S.E., Hildebrand, J.M., and Murphy, J.M. (2021). Location, location, location: A compartmentalized view of TNF-induced necroptotic signaling. Sci Signal 14. 10.1126/scisignal.abc6178.

27. Akutsu, M., Dikic, I., and Bremm, A. (2016). Ubiquitin chain diversity at a glance. J Cell Sci 129, 875–880. 10.1242/jcs.183954.

28. Komander, D., and Rape, M. (2012). The ubiquitin code. Annu Rev Biochem 81, 203–229. 10.1146/annurev-biochem-060310-170328.

29. Swatek, K.N., and Komander, D. (2016). Ubiquitin modifications. Cell Res 26, 399–422. 10.1038/cr.2016.39.

30. Gerlach, B., Cordier, S.M., Schmukle, A.C., Emmerich, C.H., Rieser, E., Haas, T.L., Webb, A.I., Rickard, J.A., Anderton, H., Wong, W.W., et al. (2011). Linear ubiquitination prevents inflammation and regulates immune signalling. Nature 471, 591–596. 10.1038/nature09816.

31. Ikeda, F., Deribe, Y.L., Skanland, S.S., Stieglitz, B., Grabbe, C., Franz-Wachtel, M., van Wijk, S.J., Goswami, P., Nagy, V., Terzic, J., et al. (2011). SHARPIN forms a linear ubiquitin ligase complex regulating NF-kappaB activity and apoptosis. Nature 471, 637–641. 10.1038/nature09814.

32. Tokunaga, F., Nakagawa, T., Nakahara, M., Saeki, Y., Taniguchi, M., Sakata, S., Tanaka, K., Nakano, H., and Iwai, K. (2011). SHARPIN is a component of the NF-kappaB-activating linear ubiquitin chain assembly complex. Nature 471, 633–636. 10.1038/nature09815.

33. Kirisako, T., Kamei, K., Murata, S., Kato, M., Fukumoto, H., Kanie, M., Sano, S., Tokunaga, F., Tanaka, K., and Iwai, K. (2006). A ubiquitin ligase complex assembles linear polyubiquitin chains. EMBO J 25, 4877–4887. 10.1038/sj.emboj.7601360.

34. Stieglitz, B., Rana, R.R., Koliopoulos, M.G., Morris-Davies, A.C., Schaeffer, V., Christodoulou, E., Howell, S., Brown, N.R., Dikic, I., and Rittinger, K. (2013). Structural basis for ligase-specific conjugation of linear ubiquitin chains by HOIP. Nature 503, 422–426. 10.1038/nature12638.

35. Smit, J.J., Monteferrario, D., Noordermeer, S.M., van Dijk, W.J., van der Reijden, B.A., and Sixma, T.K. (2012). The E3 ligase HOIP specifies linear ubiquitin chain assembly through its RING-IBR-RING domain and the unique LDD extension. EMBO J 31, 3833–3844. 10.1038/emboj.2012.217.

36. Lechtenberg, B.C., Rajput, A., Sanishvili, R., Dobaczewska, M.K., Ware, C.F., Mace, P.D., and Riedl, S.J. (2016). Structure of a HOIP/E2∼ubiquitin complex reveals RBR E3 ligase mechanism and regulation. Nature 529, 546–550. 10.1038/nature16511.

37. Oikawa, D., Sato, Y., Ohtake, F., Komakura, K., Hanada, K., Sugawara, K., Terawaki, S., Mizukami, Y., Phuong, H.T., Iio, K., et al. (2020). Molecular bases for HOIPINs-mediated inhibition of LUBAC and innate immune responses. Commun Biol 3, 163. 10.1038/s42003-020-0882-8.

38. Katsuya, K., Oikawa, D., Iio, K., Obika, S., Hori, Y., Urashima, T., Ayukawa, K., and Tokunaga, F. (2019). Small-molecule inhibitors of linear ubiquitin chain assembly complex (LUBAC), HOIPINs, suppress NF-kappaB signaling. Biochem Biophys Res Commun 509, 700–706. 10.1016/j.bbrc.2018.12.164.

39. Hrdinka, M., and Gyrd-Hansen, M. (2017). The Met1-Linked Ubiquitin Machinery: Emerging Themes of (De)regulation. Mol Cell 68, 265–280. 10.1016/j.molcel.2017.09.001.

40. Hrdinka, M., Fiil, B.K., Zucca, M., Leske, D., Bagola, K., Yabal, M., Elliott, P.R., Damgaard, R.B., Komander, D., Jost, P.J., and Gyrd-Hansen, M. (2016). CYLD Limits Lys63- and Met1-Linked Ubiquitin at Receptor Complexes to Regulate Innate Immune Signaling. Cell Rep 14, 2846–2858. 10.1016/j.celrep.2016.02.062.

41. Boisson, B., Laplantine, E., Dobbs, K., Cobat, A., Tarantino, N., Hazen, M., Lidov, H.G., Hopkins, G., Du, L., Belkadi, A., et al. (2015). Human HOIP and LUBAC deficiency underlies autoinflammation, immunodeficiency, amylopectinosis, and lymphangiectasia. J Exp Med 212, 939–951. 10.1084/jem.20141130.

42. Boisson, B., Laplantine, E., Prando, C., Giliani, S., Israelsson, E., Xu, Z., Abhyankar, A., Israel, L., Trevejo-Nunez, G., Bogunovic, D., et al. (2012). Immunodeficiency, autoinflammation and amylopectinosis in humans with inherited HOIL-1 and LUBAC deficiency. Nat Immunol 13, 1178–1186. 10.1038/ni.2457.

43. Oda, H., Manthiram, K., Chavan, P.P., Nakabo, S., Kuehn, H.S., Beck, D.B., Chae, J.J., Nehrebecky, M., Ombrello, A.K., Romeo, T., et al. (2022). Human LUBAC deficiency leads to autoinflammation and immunodeficiency by dysregulation in TNF-mediated cell death. medRxiv, 2022.2011.2009.22281431. 10.1101/2022.11.09.22281431.

44. Tokunaga, F., Sakata, S., Saeki, Y., Satomi, Y., Kirisako, T., Kamei, K., Nakagawa, T., Kato, M., Murata, S., Yamaoka, S., et al. (2009). Involvement of linear polyubiquitylation of NEMO in NF-kappaB activation. Nat Cell Biol 11, 123–132. 10.1038/ncb1821.

45. Seymour, R.E., Hasham, M.G., Cox, G.A., Shultz, L.D., Hogenesch, H., Roopenian, D.C., and Sundberg, J.P. (2007). Spontaneous mutations in the mouse Sharpin gene result in multiorgan inflammation, immune system dysregulation and dermatitis. Genes Immun 8, 416–421. 10.1038/sj.gene.6364403.

46. Rickard, J.A., Anderton, H., Etemadi, N., Nachbur, U., Darding, M., Peltzer, N., Lalaoui, N., Lawlor, K.E., Vanyai, H., Hall, C., et al. (2014). TNFR1-dependent cell death drives inflammation in Sharpin-deficient mice. Elife 3. 10.7554/eLife.03464.

47. Kumari, S., Redouane, Y., Lopez-Mosqueda, J., Shiraishi, R., Romanowska, M., Lutzmayer, S., Kuiper, J., Martinez, C., Dikic, I., Pasparakis, M., and Ikeda, F. (2014). Sharpin prevents skin inflammation by inhibiting TNFR1-induced keratinocyte apoptosis. Elife 3. 10.7554/eLife.03422.

48. Peltzer, N., Rieser, E., Taraborrelli, L., Draber, P., Darding, M., Pernaute, B., Shimizu, Y., Sarr, A., Draberova, H., Montinaro, A., et al. (2014). HOIP deficiency causes embryonic lethality by aberrant TNFR1-mediated endothelial cell death. Cell Rep 9, 153–165. 10.1016/j.celrep.2014.08.066.

49. Shimizu, Y., Peltzer, N., Sevko, A., Lafont, E., Sarr, A., Draberova, H., and Walczak, H. (2017). The Linear ubiquitin chain assembly complex acts as a liver tumor suppressor and inhibits hepatocyte apoptosis and hepatitis. Hepatology 65, 1963–1978. 10.1002/hep.29074.

50. Taraborrelli, L., Peltzer, N., Montinaro, A., Kupka, S., Rieser, E., Hartwig, T., Sarr, A., Darding, M., Draber, P., Haas, T.L., et al. (2018). LUBAC prevents lethal dermatitis by inhibiting cell death induced by TNF, TRAIL and CD95L. Nat Commun 9, 3910. 10.1038/s41467-018-06155-8.

51. Katsuya, K., Hori, Y., Oikawa, D., Yamamoto, T., Umetani, K., Urashima, T., Kinoshita, T., Ayukawa, K., Tokunaga, F., and Tamaru, M. (2018). High-Throughput Screening for Linear Ubiquitin Chain Assembly Complex (LUBAC) Selective Inhibitors Using Homogenous Time-Resolved Fluorescence (HTRF)-Based Assay System. SLAS Discov 23, 1018–1029. 10.1177/2472555218793066.

52. Shimizu, Y., Taraborrelli, L., and Walczak, H. (2015). Linear ubiquitination in immunity. Immunol Rev 266, 190–207. 10.1111/imr.12309.

53. Fan, W., Guo, J., Gao, B., Zhang, W., Ling, L., Xu, T., Pan, C., Li, L., Chen, S., Wang, H., et al. (2019). Flotillin-mediated endocytosis and ALIX-syntenin-1-mediated exocytosis protect the cell membrane from damage caused by necroptosis. Sci Signal 12. 10.1126/scisignal.aaw3423.

54. Bickel, P.E., Scherer, P.E., Schnitzer, J.E., Oh, P., Lisanti, M.P., and Lodish, H.F. (1997). Flotillin and epidermal surface antigen define a new family of caveolae-associated integral membrane proteins. J Biol Chem 272, 13793–13802. 10.1074/jbc.272.21.13793.

55. Lang, D.M., Lommel, S., Jung, M., Ankerhold, R., Petrausch, B., Laessing, U., Wiechers, M.F., Plattner, H., and Stuermer, C.A. (1998). Identification of reggie-1 and reggie-2 as plasmamembrane-associated proteins which cocluster with activated GPI-anchored cell adhesion molecules in non-caveolar micropatches in neurons. J Neurobiol 37, 502–523. 10.1002/(sici)1097-4695(199812)37:4<502::aid-neu2>3.0.co;2-s.

56. Lingwood, D., and Simons, K. (2010). Lipid rafts as a membrane-organizing principle. Science 327, 46–50. 10.1126/science.1174621.

57. Sezgin, E., Levental, I., Mayor, S., and Eggeling, C. (2017). The mystery of membrane organization: composition, regulation and roles of lipid rafts. Nat Rev Mol Cell Biol 18, 361–374. 10.1038/nrm.2017.16.

58. Morrow, I.C., Rea, S., Martin, S., Prior, I.A., Prohaska, R., Hancock, J.F., James, D.E., and Parton, R.G. (2002). Flotillin-1/reggie-2 traffics to surface raft domains via a novel golgi-independent pathway. Identification of a novel membrane targeting domain and a role for palmitoylation. J Biol Chem 277, 48834–48841. 10.1074/jbc.M209082200.

59. Solis, G.P., Hoegg, M., Munderloh, C., Schrock, Y., Malaga-Trillo, E., Rivera-Milla, E., and Stuermer, C.A. (2007). Reggie/flotillin proteins are organized into stable tetramers in membrane microdomains. Biochem J 403, 313–322. 10.1042/BJ20061686.

60. Frick, M., Bright, N.A., Riento, K., Bray, A., Merrified, C., and Nichols, B.J. (2007). Coassembly of flotillins induces formation of membrane microdomains, membrane curvature, and vesicle budding. Curr Biol 17, 1151–1156. 10.1016/j.cub.2007.05.078.

61. Glebov, O.O., Bright, N.A., and Nichols, B.J. (2006). Flotillin-1 defines a clathrin-independent endocytic pathway in mammalian cells. Nat Cell Biol 8, 46–54. 10.1038/ncb1342.

62. Langhorst, M.F., Solis, G.P., Hannbeck, S., Plattner, H., and Stuermer, C.A. (2007). Linking membrane microdomains to the cytoskeleton: regulation of the lateral mobility of reggie-1/flotillin-2 by interaction with actin. FEBS Lett 581, 4697–4703. 10.1016/j.febslet.2007.08.074.

63. Rossy, J., Schlicht, D., Engelhardt, B., and Niggli, V. (2009). Flotillins interact with PSGL-1 in neutrophils and, upon stimulation, rapidly organize into membrane domains subsequently accumulating in the uropod. PLoS One 4, e5403. 10.1371/journal.pone.0005403.

64. Affentranger, S., Martinelli, S., Hahn, J., Rossy, J., and Niggli, V. (2011). Dynamic reorganization of flotillins in chemokine-stimulated human T-lymphocytes. BMC Cell Biol 12, 28. 10.1186/1471-2121-12-28.

65. Solis, G.P., Schrock, Y., Hulsbusch, N., Wiechers, M., Plattner, H., and Stuermer, C.A. (2012). Reggies/flotillins regulate E-cadherin-mediated cell contact formation by affecting EGFR trafficking. Mol Biol Cell 23, 1812–1825. 10.1091/mbc.E11-12-1006.

66. Guillaume, E., Comunale, F., Do Khoa, N., Planchon, D., Bodin, S., and Gauthier-Rouviere, C. (2013). Flotillin microdomains stabilize cadherins at cell-cell junctions. J Cell Sci 126, 5293–5304. 10.1242/jcs.133975.

67. Amaddii, M., Meister, M., Banning, A., Tomasovic, A., Mooz, J., Rajalingam, K., and Tikkanen, R. (2012). Flotillin-1/reggie-2 protein plays dual role in activation of receptor-tyrosine kinase/mitogen-activated protein kinase signaling. J Biol Chem 287, 7265–7278. 10.1074/jbc.M111.287599.

68. Pust, S., Klokk, T.I., Musa, N., Jenstad, M., Risberg, B., Erikstein, B., Tcatchoff, L., Liestol, K., Danielsen, H.E., van Deurs, B., and Sandvig, K. (2013). Flotillins as regulators of ErbB2 levels in breast cancer. Oncogene 32, 3443–3451. 10.1038/onc.2012.357.

69. Damgaard, R.B., Elliott, P.R., Swatek, K.N., Maher, E.R., Stepensky, P., Elpeleg, O., Komander, D., and Berkun, Y. (2019). OTULIN deficiency in ORAS causes cell type-specific LUBAC degradation, dysregulated TNF signalling and cell death. EMBO Mol Med 11, e9324. 10.15252/emmm.201809324.

70. Damgaard, R.B., Walker, J.A., Marco-Casanova, P., Morgan, N.V., Titheradge, H.L., Elliott, P.R., McHale, D., Maher, E.R., McKenzie, A.N.J., and Komander, D. (2016). The Deubiquitinase OTULIN Is an Essential Negative Regulator of Inflammation and Autoimmunity. Cell 166, 1215–1230 e1220. 10.1016/j.cell.2016.07.019.

71. Sakamoto, H., Egashira, S., Saito, N., Kirisako, T., Miller, S., Sasaki, Y., Matsumoto, T., Shimonishi, M., Komatsu, T., Terai, T., et al. (2015). Gliotoxin suppresses NF-kappaB activation by selectively inhibiting linear ubiquitin chain assembly complex (LUBAC). ACS Chem Biol 10, 675–681. 10.1021/cb500653y.

72. Peltzer, N., Darding, M., Montinaro, A., Draber, P., Draberova, H., Kupka, S., Rieser, E., Fisher, A., Hutchinson, C., Taraborrelli, L., et al. (2018). LUBAC is essential for embryogenesis by preventing cell death and enabling haematopoiesis. Nature 557, 112–117. 10.1038/s41586-018-0064-8.

73. Vanlangenakker, N., Bertrand, M.J., Bogaert, P., Vandenabeele, P., and Vanden Berghe, T. (2011). TNF-induced necroptosis in L929 cells is tightly regulated by multiple TNFR1 complex I and II members. Cell Death Dis 2, e230. 10.1038/cddis.2011.111.

74. Zargarian, S., Shlomovitz, I., Erlich, Z., Hourizadeh, A., Ofir-Birin, Y., Croker, B.A., Regev-Rudzki, N., Edry-Botzer, L., and Gerlic, M. (2017). Phosphatidylserine externalization, “necroptotic bodies” release, and phagocytosis during necroptosis. PLoS Biol 15, e2002711. 10.1371/journal.pbio.2002711.

75. Yoon, S., Kovalenko, A., Bogdanov, K., and Wallach, D. (2017). MLKL, the Protein that Mediates Necroptosis, Also Regulates Endosomal Trafficking and Extracellular Vesicle Generation. Immunity 47, 51–65 e57. 10.1016/j.immuni.2017.06.001.

76. Eiraku, M., and Sasai, Y. (2012). Self-formation of layered neural structures in three-dimensional culture of ES cells. Curr Opin Neurobiol 22, 768–777. 10.1016/j.conb.2012.02.005.

77. Lancaster, M.A., and Knoblich, J.A. (2014). Organogenesis in a dish: modeling development and disease using organoid technologies. Science 345, 1247125. 10.1126/science.1247125.

78. Fatehullah, A., Tan, S.H., and Barker, N. (2016). Organoids as an in vitro model of human development and disease. Nat Cell Biol 18, 246–254. 10.1038/ncb3312.

79. Boj, S.F., Hwang, C.I., Baker, L.A., Chio, II, Engle, D.D., Corbo, V., Jager, M., Ponz-Sarvise, M., Tiriac, H., Spector, M.S., et al. (2015). Organoid models of human and mouse ductal pancreatic cancer. Cell 160, 324–338. 10.1016/j.cell.2014.12.021.

80. Broutier, L., Andersson-Rolf, A., Hindley, C.J., Boj, S.F., Clevers, H., Koo, B.K., and Huch, M. (2016). Culture and establishment of self-renewing human and mouse adult liver and pancreas 3D organoids and their genetic manipulation. Nat Protoc 11, 1724–1743. 10.1038/nprot.2016.097.

81. Dossena, M., Piras, R., Cherubini, A., Barilani, M., Dugnani, E., Salanitro, F., Moreth, T., Pampaloni, F., Piemonti, L., and Lazzari, L. (2020). Standardized GMP-compliant scalable production of human pancreas organoids. Stem Cell Res Ther 11, 94. 10.1186/s13287-020-1585-2.

82. Georgakopoulos, N., Prior, N., Angres, B., Mastrogiovanni, G., Cagan, A., Harrison, D., Hindley, C.J., Arnes-Benito, R., Liau, S.S., Curd, A., et al. (2020). Long-term expansion, genomic stability and in vivo safety of adult human pancreas organoids. BMC Dev Biol 20, 4. 10.1186/s12861-020-0209-5.

83. Jung, N., Moreth, T., Stelzer, E.H.K., Pampaloni, F., and Windbergs, M. (2021). Non-invasive analysis of pancreas organoids in synthetic hydrogels defines material-cell interactions and luminal composition. Biomater Sci 9, 5415–5426. 10.1039/d1bm00597a.

84. Mocarski, E.S., Guo, H., and Kaiser, W.J. (2015). Necroptosis: The Trojan horse in cell autonomous antiviral host defense. Virology 479-480, 160-166. 10.1016/j.virol.2015.03.016.

85. Kaiser, W.J., Upton, J.W., and Mocarski, E.S. (2013). Viral modulation of programmed necrosis. Curr Opin Virol 3, 296–306. 10.1016/j.coviro.2013.05.019.

86. Pearson, J.S., Giogha, C., Muhlen, S., Nachbur, U., Pham, C.L., Zhang, Y., Hildebrand, J.M., Oates, C.V., Lung, T.W., Ingle, D., et al. (2017). EspL is a bacterial cysteine protease effector that cleaves RHIM proteins to block necroptosis and inflammation. Nat Microbiol 2, 16258. 10.1038/nmicrobiol.2016.258.

87. Upton, J.W., and Kaiser, W.J. (2017). DAI Another Way: Necroptotic Control of Viral Infection. Cell Host Microbe 21, 290–293. 10.1016/j.chom.2017.01.016.

88. Seifert, L., Werba, G., Tiwari, S., Giao Ly, N.N., Alothman, S., Alqunaibit, D., Avanzi, A., Barilla, R., Daley, D., Greco, S.H., et al. (2016). The necrosome promotes pancreatic oncogenesis via CXCL1 and Mincle-induced immune suppression. Nature 532, 245–249. 10.1038/nature17403.

89. Najafov, A., Chen, H., and Yuan, J. (2017). Necroptosis and Cancer. Trends Cancer 3, 294–301. 10.1016/j.trecan.2017.03.002.

90. Pefanis, A., Ierino, F.L., Murphy, J.M., and Cowan, P.J. (2019). Regulated necrosis in kidney ischemia-reperfusion injury. Kidney Int 96, 291–301. 10.1016/j.kint.2019.02.009.

91. Muller, T., Dewitz, C., Schmitz, J., Schroder, A.S., Brasen, J.H., Stockwell, B.R., Murphy, J.M., Kunzendorf, U., and Krautwald, S. (2017). Necroptosis and ferroptosis are alternative cell death pathways that operate in acute kidney failure. Cell Mol Life Sci 74, 3631–3645. 10.1007/s00018-017-2547-4.

92. Petrie, E.J., Sandow, J.J., Jacobsen, A.V., Smith, B.J., Griffin, M.D.W., Lucet, I.S., Dai, W., Young, S.N., Tanzer, M.C., Wardak, A., et al. (2018). Conformational switching of the pseudokinase domain promotes human MLKL tetramerization and cell death by necroptosis. Nat Commun 9, 2422. 10.1038/s41467-018-04714-7.

93. Davies, K.A., Tanzer, M.C., Griffin, M.D.W., Mok, Y.F., Young, S.N., Qin, R., Petrie, E.J., Czabotar, P.E., Silke, J., and Murphy, J.M. (2018). The brace helices of MLKL mediate interdomain communication and oligomerisation to regulate cell death by necroptosis. Cell Death Differ 25, 1567–1580. 10.1038/s41418-018-0061-3.

94. Tanzer, M.C., Matti, I., Hildebrand, J.M., Young, S.N., Wardak, A., Tripaydonis, A., Petrie, E.J., Mildenhall, A.L., Vaux, D.L., Vince, J.E., et al. (2016). Evolutionary divergence of the necroptosis effector MLKL. Cell Death Differ 23, 1185–1197. 10.1038/cdd.2015.169.

95. Petrie, E.J., Czabotar, P.E., and Murphy, J.M. (2019). The Structural Basis of Necroptotic Cell Death Signaling. Trends Biochem Sci 44, 53–63. 10.1016/j.tibs.2018.11.002.

96. Tanzer, M.C., Tripaydonis, A., Webb, A.I., Young, S.N., Varghese, L.N., Hall, C., Alexander, W.S., Hildebrand, J.M., Silke, J., and Murphy, J.M. (2015). Necroptosis signalling is tuned by phosphorylation of MLKL residues outside the pseudokinase domain activation loop. Biochem J 471, 255–265. 10.1042/BJ20150678.

97. Rodriguez, D.A., Weinlich, R., Brown, S., Guy, C., Fitzgerald, P., Dillon, C.P., Oberst, A., Quarato, G., Low, J., Cripps, J.G., et al. (2016). Characterization of RIPK3-mediated phosphorylation of the activation loop of MLKL during necroptosis. Cell Death Differ 23, 76–88. 10.1038/cdd.2015.70.

98. Dai, J., Zhang, C., Guo, L., He, H., Jiang, K., Huang, Y., Zhang, X., Zhang, H., Wei, W., Zhang, Y., et al. (2020). A necroptotic-independent function of MLKL in regulating endothelial cell adhesion molecule expression. Cell Death Dis 11, 282. 10.1038/s41419-020-2483-3.

99. Petrie, E.J., Birkinshaw, R.W., Koide, A., Denbaum, E., Hildebrand, J.M., Garnish, S.E., Davies, K.A., Sandow, J.J., Samson, A.L., Gavin, X., et al. (2020). Identification of MLKL membrane translocation as a checkpoint in necroptotic cell death using Monobodies. Proc Natl Acad Sci U S A 117, 8468–8475. 10.1073/pnas.1919960117.

100. Douanne, T., Andre-Gregoire, G., Trillet, K., Thys, A., Papin, A., Feyeux, M., Hulin, P., Chiron, D., Gavard, J., and Bidere, N. (2019). Pannexin-1 limits the production of proinflammatory cytokines during necroptosis. EMBO Rep 20, e47840. 10.15252/embr.201947840.

101. Mizuno, E., Kawahata, K., Kato, M., Kitamura, N., and Komada, M. (2003). STAM proteins bind ubiquitinated proteins on the early endosome via the VHS domain and ubiquitin-interacting motif. Mol Biol Cell 14, 3675–3689. 10.1091/mbc.e02-12-0823.

102. Lohi, O., and Lehto, V.P. (1998). VHS domain marks a group of proteins involved in endocytosis and vesicular trafficking. FEBS Lett 440, 255–257. 10.1016/s0014-5793(98)01401-x.

103. Hirano, S., Kawasaki, M., Ura, H., Kato, R., Raiborg, C., Stenmark, H., and Wakatsuki, S. (2006). Double-sided ubiquitin binding of Hrs-UIM in endosomal protein sorting. Nat Struct Mol Biol 13, 272–277. 10.1038/nsmb1051.

104. Bache, K.G., Raiborg, C., Mehlum, A., and Stenmark, H. (2003). STAM and Hrs are subunits of a multivalent ubiquitin-binding complex on early endosomes. J Biol Chem 278, 12513–12521. 10.1074/jbc.M210843200.

105. Pornillos, O., Alam, S.L., Rich, R.L., Myszka, D.G., Davis, D.R., and Sundquist, W.I. (2002). Structure and functional interactions of the Tsg101 UEV domain. EMBO J 21, 2397–2406. 10.1093/emboj/21.10.2397.

106. Sundquist, W.I., Schubert, H.L., Kelly, B.N., Hill, G.C., Holton, J.M., and Hill, C.P. (2004). Ubiquitin recognition by the human TSG101 protein. Mol Cell 13, 783–789. 10.1016/s1097-2765(04)00129-7.

107. Teo, H., Veprintsev, D.B., and Williams, R.L. (2004). Structural insights into endosomal sorting complex required for transport (ESCRT-I) recognition of ubiquitinated proteins. J Biol Chem 279, 28689–28696. 10.1074/jbc.M400023200.

108. Meister, M., Banfer, S., Gartner, U., Koskimies, J., Amaddii, M., Jacob, R., and Tikkanen, R. (2017). Regulation of cargo transfer between ESCRT-0 and ESCRT-I complexes by flotillin-1 during endosomal sorting of ubiquitinated cargo. Oncogenesis 6, e344. 10.1038/oncsis.2017.47.

109. Legler, D.F., Micheau, O., Doucey, M.A., Tschopp, J., and Bron, C. (2003). Recruitment of TNF receptor 1 to lipid rafts is essential for TNFalpha-mediated NF-kappaB activation. Immunity 18, 655–664. 10.1016/s1074-7613(03)00092-x.

110. Hunter, I., and Nixon, G.F. (2006). Spatial compartmentalization of tumor necrosis factor (TNF) receptor 1-dependent signaling pathways in human airway smooth muscle cells. Lipid rafts are essential for TNF-alpha-mediated activation of RhoA but dispensable for the activation of the NF-kappaB and MAPK pathways. J Biol Chem 281, 34705–34715. 10.1074/jbc.M605738200.

111. Maminska, A., Bartosik, A., Banach-Orlowska, M., Pilecka, I., Jastrzebski, K., Zdzalik-Bielecka, D., Castanon, I., Poulain, M., Neyen, C., Wolinska-Niziol, L., et al. (2016). ESCRT proteins restrict constitutive NF-kappaB signaling by trafficking cytokine receptors. Sci Signal 9, ra8. 10.1126/scisignal.aad0848.

112. Mahul-Mellier, A.L., Strappazzon, F., Petiot, A., Chatellard-Causse, C., Torch, S., Blot, B., Freeman, K., Kuhn, L., Garin, J., Verna, J.M., et al. (2008). Alix and ALG-2 are involved in tumor necrosis factor receptor 1-induced cell death. J Biol Chem 283, 34954–34965. 10.1074/jbc.M803140200.

113. Liu, Z., Dagley, L.F., Shield-Artin, K., Young, S.N., Bankovacki, A., Wang, X., Tang, M., Howitt, J., Stafford, C.A., Nachbur, U., et al. (2021). Oligomerization-driven MLKL ubiquitylation antagonises necroptosis. bioRxiv, 2021.2005.2001.442209. 10.1101/2021.05.01.442209.

114. Garcia, L.R., Tenev, T., Newman, R., Haich, R.O., Liccardi, G., John, S.W., Annibaldi, A., Yu, L., Pardo, M., Young, S.N., et al. (2021). Ubiquitylation of MLKL at lysine 219 positively regulates necroptosis-induced tissue injury and pathogen clearance. Nat Commun 12, 3364. 10.1038/s41467-021-23474-5.

115. Akimov, V., Barrio-Hernandez, I., Hansen, S.V.F., Hallenborg, P., Pedersen, A.K., Bekker-Jensen, D.B., Puglia, M., Christensen, S.D.K., Vanselow, J.T., Nielsen, M.M., et al. (2018). UbiSite approach for comprehensive mapping of lysine and N-terminal ubiquitination sites. Nat Struct Mol Biol 25, 631–640. 10.1038/s41594-018-0084-y.

116. Roedig, J., Kowald, L., Juretschke, T., Karlowitz, R., Ahangarian Abhari, B., Roedig, H., Fulda, S., Beli, P., and van Wijk, S.J. (2021). USP22 controls necroptosis by regulating receptor-interacting protein kinase 3 ubiquitination. EMBO Rep 22, e50163. 10.15252/embr.202050163.

117. Muller, S., Rycak, L., Afonso-Grunz, F., Winter, P., Zawada, A.M., Damrath, E., Scheider, J., Schmah, J., Koch, I., Kahl, G., and Rotter, B. (2014). APADB: a database for alternative polyadenylation and microRNA regulation events. Database (Oxford) 2014. 10.1093/database/bau076.

118. Martin, M. (2011). Cutadapt removes adapter sequences from high-throughput sequencing reads. 2011 17, 3. 10.14806/ej.17.1.200.

119. Love, M.I., Huber, W., and Anders, S. (2014). Moderated estimation of fold change and dispersion for RNA-seq data with DESeq2. Genome Biol 15, 550. 10.1186/s13059-014-0550-8.

