## Supplementary figures and images for "Species-specific LUBAC-mediated M1 ubiquitination regulates necroptosis by segregating the cellular distribution and fate of activated MLKL"

### Supplemental Information

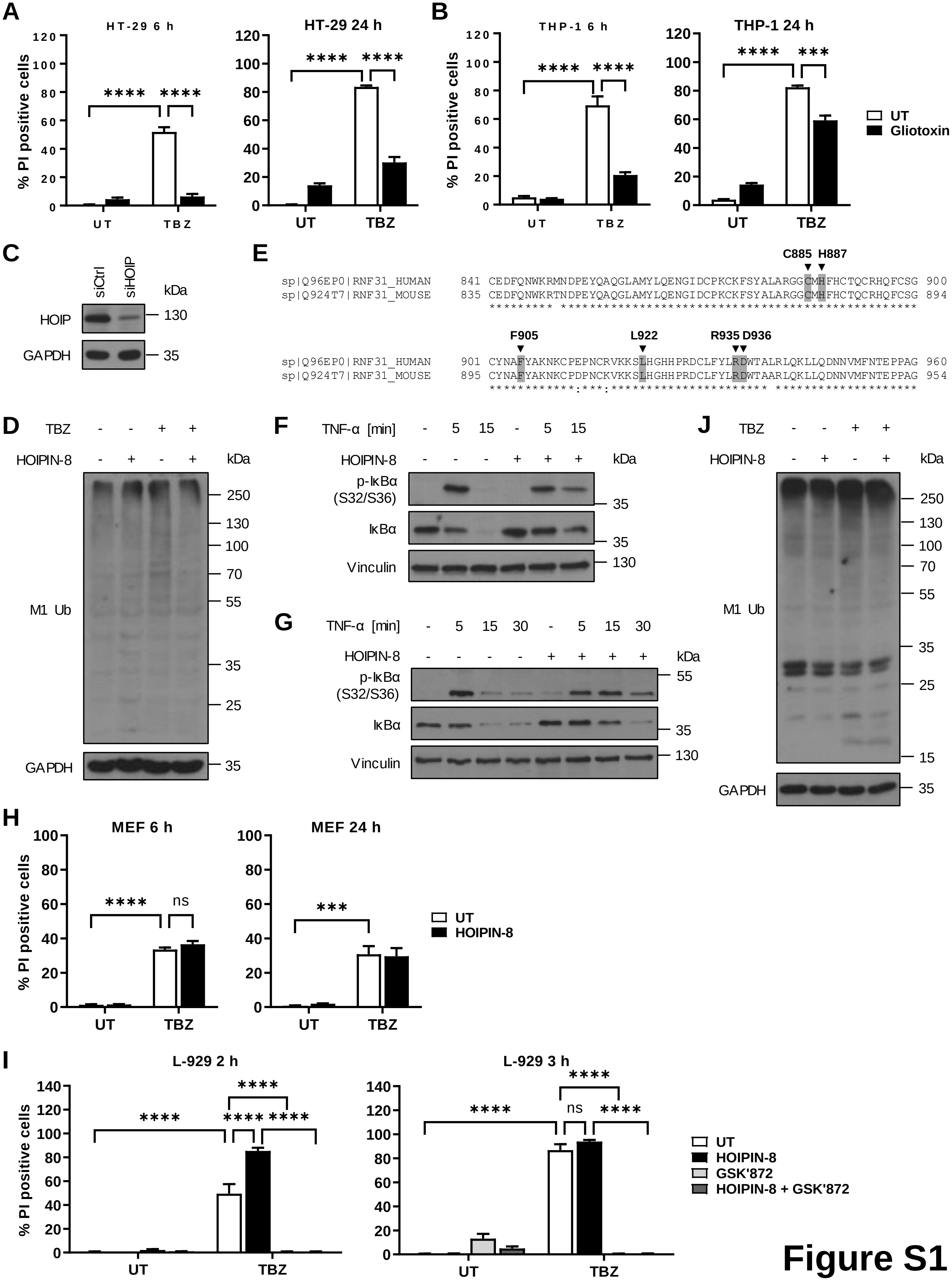

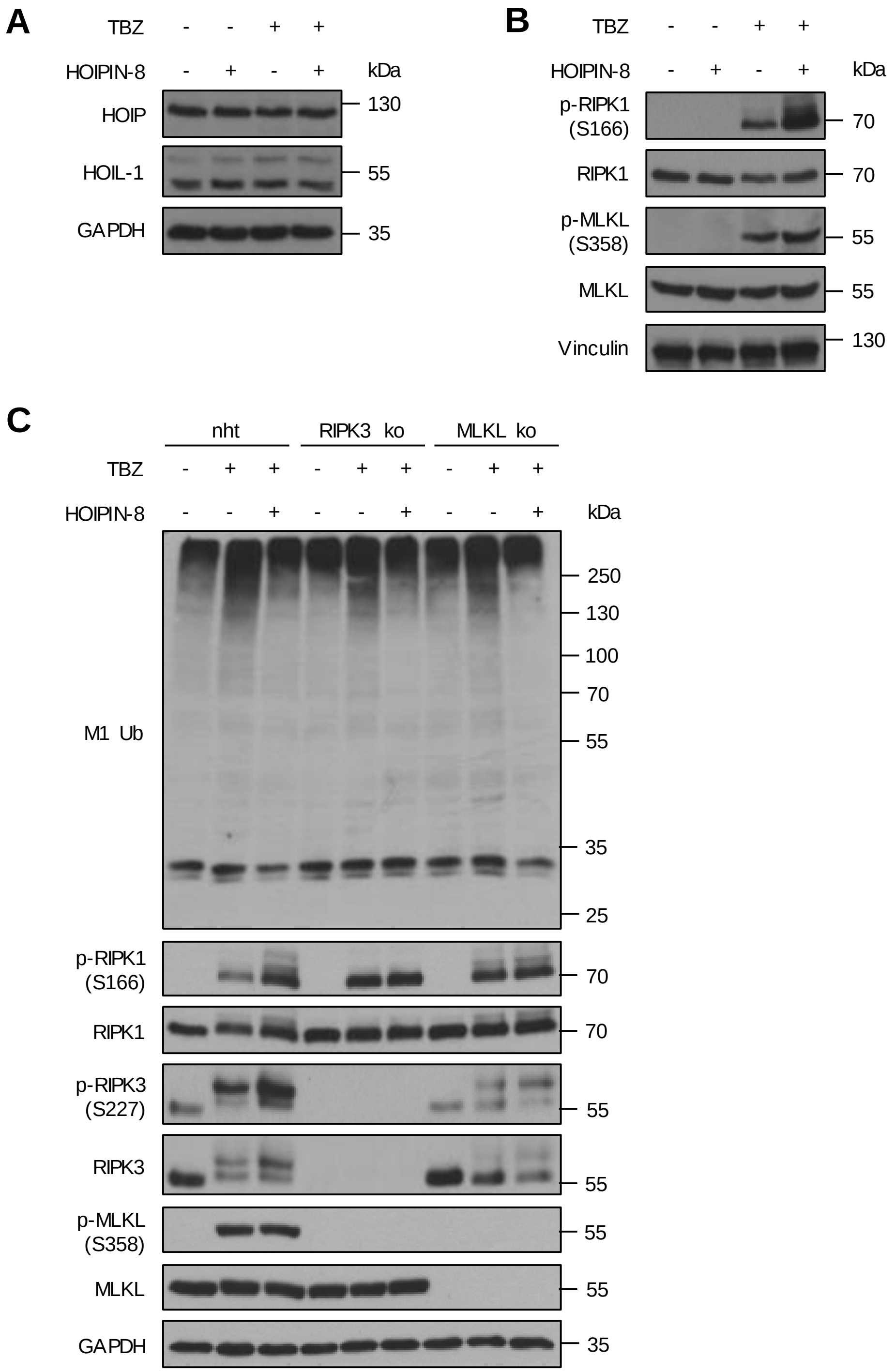

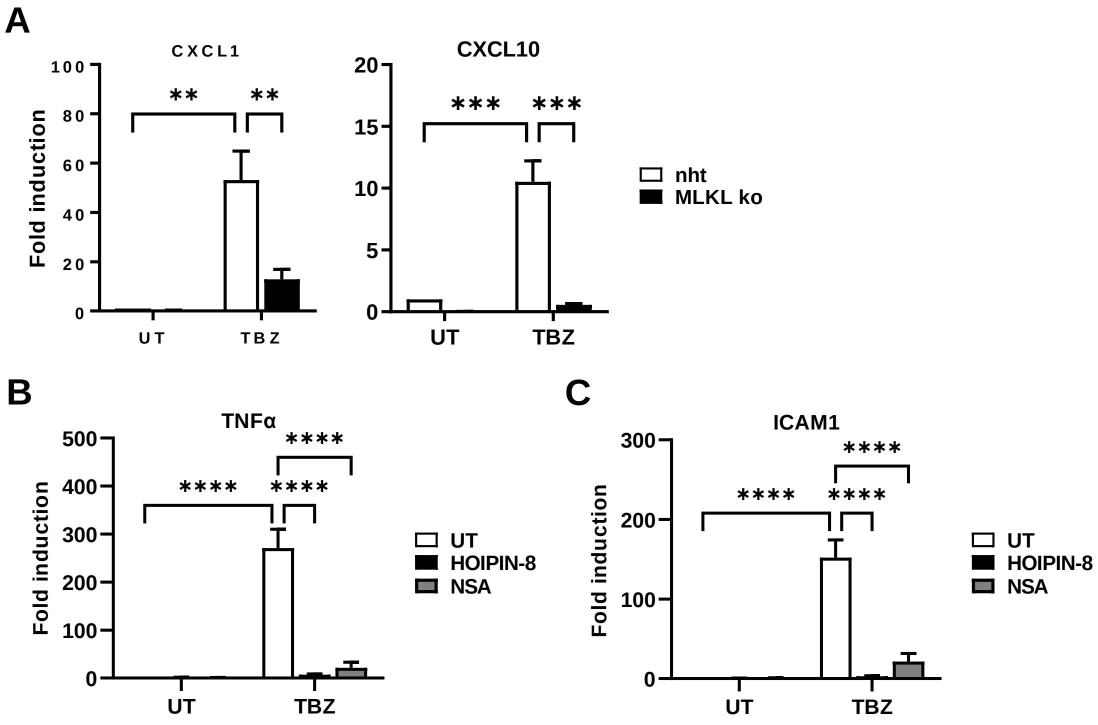

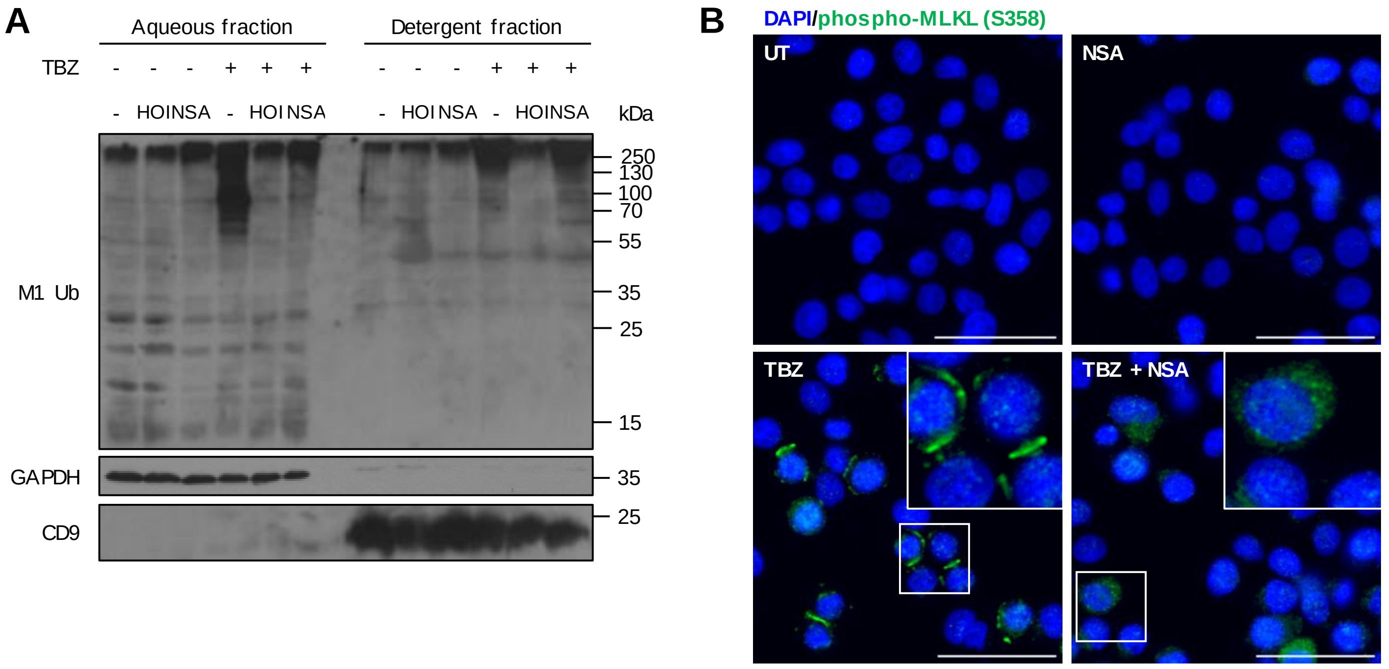

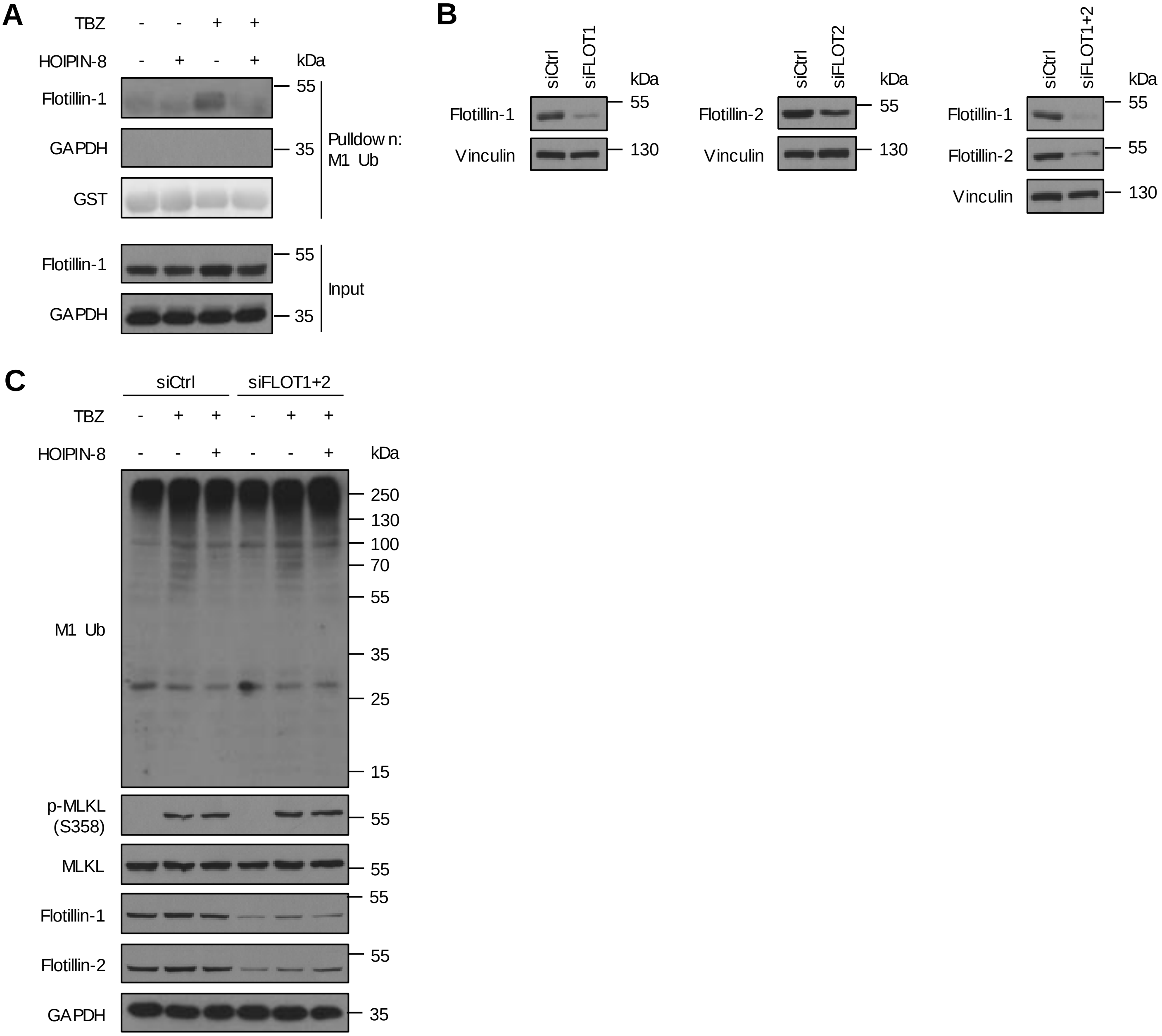

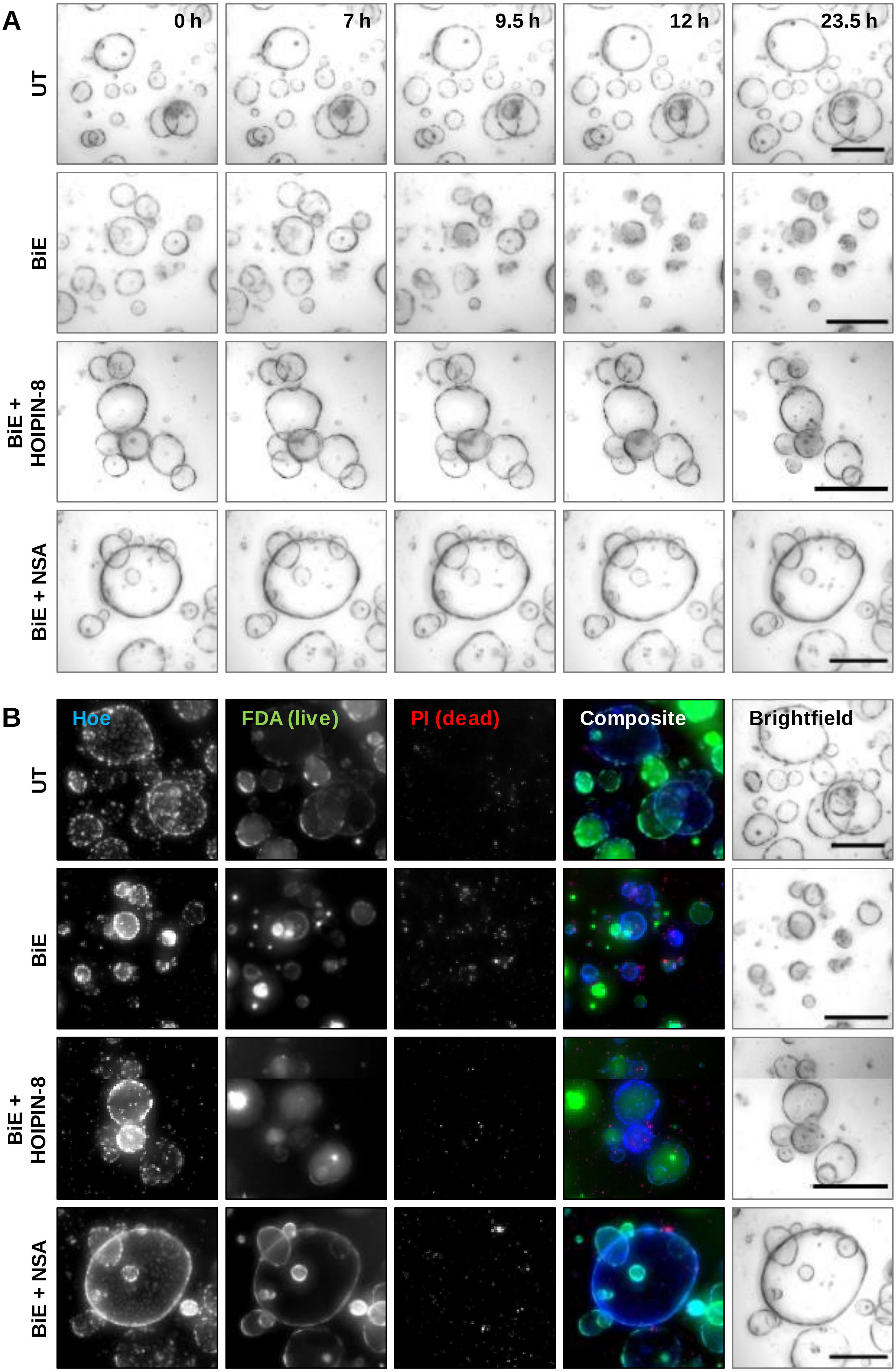

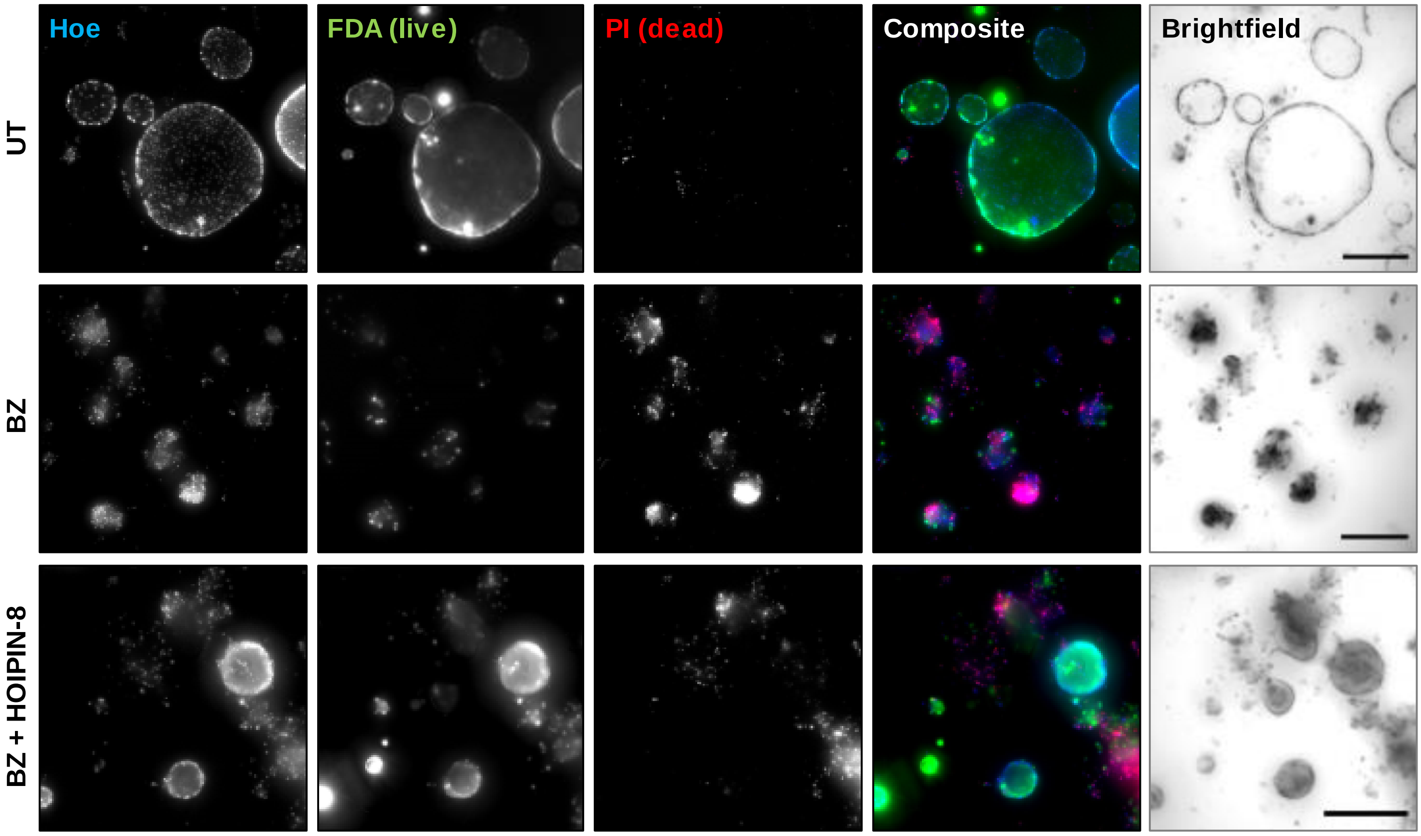


#### Figure S7
